# Distinct brain mechanisms support trust violations, belief integration, and bias in human-AI teams

**DOI:** 10.1101/2025.08.16.670645

**Authors:** Luisa Roeder, Pamela Hoyte, Graham Kerr, Peter Bruza, Johan N van der Meer

**Author notes:** **For correspondence:** (LR); (PB); (JN). Faculty of Science, Queensland University of Technology, Brisbane, Australia (Peter Bruza).

## Abstract

This study provides an integrated electrophysiological and behavioral account of the neuro-cognitive markers underlying trust evolution during human interaction with artificial intelligence (AI). Trust is essential for effective collaboration and plays a key role in realizing the benefits of human–AI teaming in information-rich and decision-critical contexts. Using electroencephalography (EEG), we identified neural signatures of dynamic shifts in human trust during a face classification task involving an AI agent. Viewing the AI’s classification elicited an N2–P3a–P3b event-related potential (ERP) complex that was sensitive to agreement with the participant’s own judgment and modulated by individual response biases. In addition, we observed a centro-parietal positivity (CPP) prior to participants’ responses, and found that ongoing EEG activity in this time window co-varied with subsequent changes in AI trust ratings. These neural effects showed substantial individual variability, indicating the use of diverse metacognitive strategies. Together, these findings suggest that trust in AI is shaped by internal confidence signals and evaluative processing of feedback.

## Introduction

Human–AI teaming is becoming increasingly common in high-stakes domains such as disaster relief, medical diagnostics, and autonomous systems. In these settings, effective collaboration depends not only on the AI agent’s ability to act reliably and competently, but also on the human partner’s capacity to form and adapt trust. Trust in this context entails a willingness to rely on another entity under uncertainty, accepting potential vulnerability in exchange for achieving shared goals (***Simpson, 2007***). A critical factor in this process is perceived reliability (***Mayer et al. (1995***), how consistently the AI performs as expected, how frequently it violates those expectations), which we adopt here as a measurable proxy for trust.

Recent electroencephalography (EEG) studies have begun to reveal how the brain monitors AI behavior in cooperative contexts (***De Visser et al., 2018***; ***Somon et al., 2019***). When an AI agent makes an error during a task with clear correct responses, observers show neural signatures of performance monitoring; in particular, the observational error-related negativity (oERN) and error positivity (oPe), which are two well-established event-related potentials (ERPs) associated with detecting and evaluating others’ mistakes (***van Schie et al., 2004***; ***Carp et al., 2009***; ***Koban et al., 2010***; ***Koban and Pourtois, 2014***; ***de Bruijn and von Rhein, 2012***). These signals suggest that the human brain tracks AI decisions using neural computations similar to that used in social or supervisory contexts.

However, these studies typically rely on tasks with an objective ground truth, where errors are unambiguous and outcomes are known (e.g., correct/incorrect responses in flanker tasks). In real-world settings, collaboration with AI agents often takes place under conditions of uncertainty, where feedback is ambiguous or unavailable, and trust must be constructed and adjusted based on internal inference (***Freedy et al***., 2007; ***Haring et al***., 2021). In such cases, we still lack a mechanistic understanding of how neural systems support belief formation, expectancy violations, or trust adaptations during ongoing interaction with an AI partner.

To address this gap, we investigated the neural dynamics of trust formation in a cooperative human–AI task with ambiguous feedback. Participants engaged in a face classification task in which they judged if images were real or AI-generated while we recorded EEG. Although the AI’s responses were pre-programmed to disagree with the participant on a fixed proportion of trials (25%), participants were briefed to believe that the (simulated) AI was making an independent, parallel judgment and that the AI classification may not necessarily be correct. In each trial, the participant responded first, followed by a brief delay, after which the AI’s classification was revealed. This sequential structure was designed to evoke a sense of cooperative interaction and agency attribution, allowing us to examine not only how participants processed the AI’s responses, but also how they formed their own decisions, potentially influencing downstream judgments of trust. A key feature of the design was the inclusion of regular behavioral measures of trust using a continuous visual analogue scale (VAS) through which participants rated the AI’s reliability throughout the task. This enabled us to track evolving impressions of trust across the entire session. This design allowed us to dissociate neural signals related to self-generated decision-making, agreement or disagreement with the AI, and changes in perceived trust over time.

Traditional trust paradigms often rely on fixed intervals (with a possible variable inter-trial interval) between decision and outcome, for example, in economic risk tasks where a high- or low-stakes wager is followed by a clearly defined gain or loss (***Wang et al***., 2016; ***Fu et al***., 2019; ***Li et al***., 2025; ***Krueger et al***., 2007). While these designs are well controlled, they may lack ecological validity and constrain the analysis of temporally precise neural dynamics. In contrast, our paradigm was structured to allow immediate EEG time-locking to the participant’s decision, enabling analysis of response-locked signals such as the centro-parietal positivity (CPP), which reflects internal decision formation. The AI classification was presented shortly afterward, allowing us to also time-lock the EEG to the feedback (in this article we refer to the event when the AI classification is revealed as ‘AI feedback’ as this is a common term in ERP research; in this study, the AI feedback is not referring to feedback on correct or incorrect performance of the human), and to capture neural responses to agreement as well as violations of expectation arising from AI disagreement under uncertainty.

Beyond condition-specific ERPs, we also explored whether any EEG signals co-varied with changes in perceived AI reliability, to identify neural activity that tracked updates in trust. Although we did not target a specific component or latency in advance, we focused on temporally precise EEG signals that might reflect internally generated processes relevant to trust formation, including both early decision-related activity and later responses to AI behavior. Finally, we examined whether individual differences in response bias (e.g., a tendency to judge more faces as fake or real) influenced how AI disagreement and agreement were processed. Such effects could reveal how personal internal strategies shape the interpretation of feedback and contribute to trust dynamics.

Together, these analyses allowed us to dissociate multiple neural processes that contribute to trust evolution during human–AI interaction: immediate neural responses to agreement or violation following AI classification; internally driven signals related to the formation of participants’ own judgments; EEG signals that co-varied with changes in perceived AI reliability; and inter-individual biases that influenced how AI classifications were processed. By examining these components in parallel, we aim to provide a mechanistic account of how trust develops and fluctuates during ongoing interaction with an AI partner.

## Results

Participants (n = 34; mean age = 26 years, 18 female) completed 560 trials of a face classification task while EEG was recorded from 64 scalp electrodes. In each trial, they judged whether a face was real or AI-generated, followed by the AI’s classification, which was pre-programmed to agree with the participant for 75% of trials. This yielded four possible human–AI response conditions: HRAIR (Human-Real & AI-Real), HFAIF (Human-Fake & AI-Fake), HRAIF (Human-Real & AI-Fake), HFAIR (Human-Fake & AI-Real). Participants rated the AI’s perceived reliability on a continuous scale at regular intervals, and behavioral measures such as response times were also collected.

We computed event-related potentials (ERPs) time-locked to the participant’s response (image judgment), capturing both neural activity related to self-generated decision-making and subsequent processing of the AI’s classification, which was presented 300 ms later. To capture the variability associated with different task conditions, we estimated single-subject beta parameter maps using the LIMO EEG toolbox (***Pernet et al***., 2011), effectively yielding condition-specific general linear model (GLM) estimates of ERP responses. EEG data were preprocessed and cleaned, resulting in an average of 81% usable trials per participant; the same cleaned trials were used for behavioral analyses.

### Behavioral results

Participants responded within the 2-second deadline on nearly all trials (missed responses <1%). Overall classification accuracy was near chance (50% ± 6%), consistent with the ambiguous nature of the stimuli. Most participants showed a preference for judging images as either fake or real, which we interpret as a response bias (Figure 1A). To explore whether this bias influenced behavior, we conducted a follow-up analysis of response times (two-way ANOVA with factors bias and response time). Participants were divided into two groups (bias towards real: n = 21; bias towards fake: n = 13), and we found a significant interaction: participants were faster when making their preferred response (e.g., those biased toward “real” were faster in real trials), F(1, 32) = 21.94, p < 0.001). Although modest in effect size (92 ms), this effect indicates consistent task engagement and the use of individualized decision strategies (Figure 1B).

**Figure 1.**
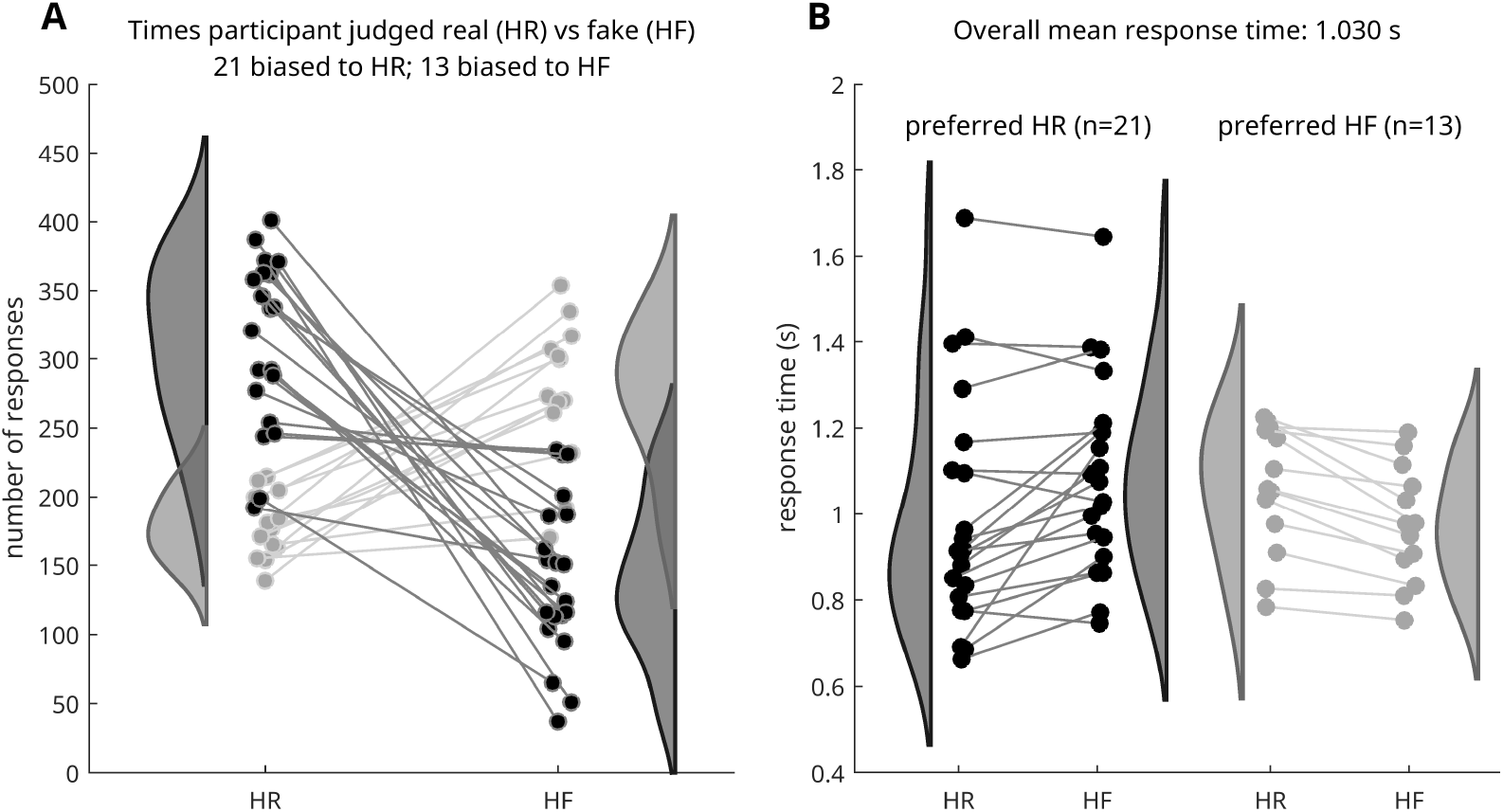
Behavioral results. **A**. Total number of responses (image judgments) if face is real (HR) and if face is fake (HF) for each of the 34 participants. Two subgroups for preference emerged for pressing real (21 participants; black dots and lines) and pressing fake (13 participants; gray dots and lines). Distributions of responses of the two subgroups are shown on the sides of the panel (dark gray shaded area, preference real; light gray shaded area, preference fake). **B**. Mean response times per participant of both groups for real (HR) and fake (HF) image judgments; participants were distributed in terms of their overall response time, with a faster response time for the most frequent judgment. Distributions (gray shaded areas) of response times are shown next to individual responses. HF, Human Fake; HR, Human Real.

### EEG markers of decision-making and feedback integration

To provide an overview of the temporal dynamics, we computed grand-average ERPs (20% trimmed mean) for each of the four human–AI response conditions: HRAIR (Human-Real & AI-Real), HFAIF (Human-Fake & AI-Fake), HRAIF (Human-Real & AI-Fake), and HFAIR (Human-Fake & AI-Real), timelocked to the onset of the AI classification (Figure 2). Baseline correction was applied using the fixation cross period preceding stimulus presentation. This choice preserves the positive deflections starting at −650 ms observed consistently across all conditions, rather than removing them.

**Figure 2.**
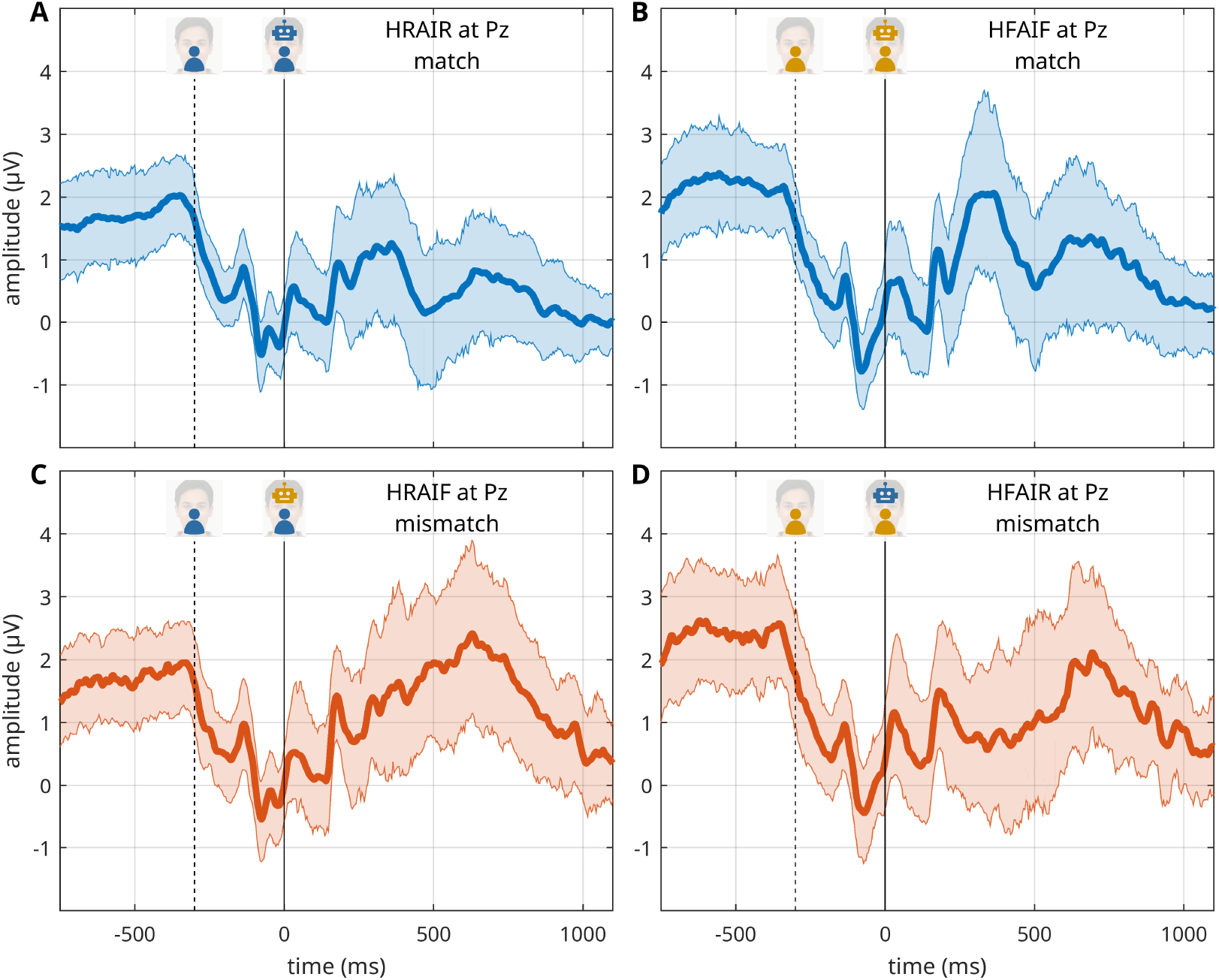
Grand average event-related potentials (ERPs). Time series of ERPs at Pz time-locked to the event when the AI classification icon (robot icon) is displayed on screen (time 0 ms) in the mismatch (red) and match (blue) conditions. At −300 ms participants made their judgment (fake or real key press). The match condition contains HRAIR and HFAIF, the mismatch condition contains HRAIF and HFAIR. The mean ERP trace is computed as the 20% trimmed mean of the mean participant-level ERP data, shaded areas denote the 95% Bayesian Highest Density Interval (see text for details). HRAIR, Human-Real & AI-Real; HFAIF, Human-Fake & AI-Fake; HRAIF, Human-Real & AI-Fake; HFAIR, Human-Fake & AI-Real.

Around 150 ms (i.e., 150 ms after the participant’s response), all conditions exhibit a brief positive deflection, likely reflecting visual processing of the human judgment icon shown after each decision. Following the onset of the AI classification at 0 ms, ERP waveforms display a series of condition-sensitive deflections. Notably, mismatch conditions (Figure 2C and D) appear to elicit a more sustained positivity from 400 ms onward compared to match conditions (Figure 2A and B).

To formally test these effects, we used a hierarchical linear modeling approach implemented in the LIMO EEG toolbox (***Pernet et al***., 2011), which estimates condition-specific beta parameters for each participant. These beta estimates reflect the contribution of each condition to the EEG signal, time-locked to the AI classification, while controlling for shared variance across conditions. Unlike the descriptive grand-average waveforms shown in Figure 2, these beta weights are used as the basis for statistical contrasts between conditions. This way, the beta estimates provide a model-based decomposition of the ERP signal, isolating variance uniquely attributable to each experimental condition (see supplementary Figure S2 for grand-average betas).

#### Neural responses to agreement and disagreement with AI

To examine how the brain processes agreement versus disagreement with the AI, we conducted a 2 × 2 repeated-measures ANOVA with the factors: (1) AI classification type (mismatch vs. match), and (2) human judgment of the image (fake vs. real). The analysis used single-subject beta estimates derived from the four experimental conditions described in the previous section. A significant main effect of AI classification (mismatch vs. match) was observed, with one large spatio-temporal cluster spanning all 64 electrodes and the time interval from –176 to 1024 ms (average F = 12.71, average p = 0.01; Figure 3A). To better isolate temporally distinct ERP components within this broad window, we conducted post hoc t-tests on the mismatch-minus-match contrast using a more stringent cluster-forming threshold (*α* = 0.01; supplementary Figure S3). This analysis revealed three significant time windows: an early negative deflection consistent with the N2 component (200–360 ms, peak at 292 ms), a mid-latency positivity consistent with P3a (404–536 ms, peak at 460 ms), and a later centro-parietal positivity consistent with P3b (532–752 ms, peak at 700 ms) (Figure 3B).

**Figure 3.**
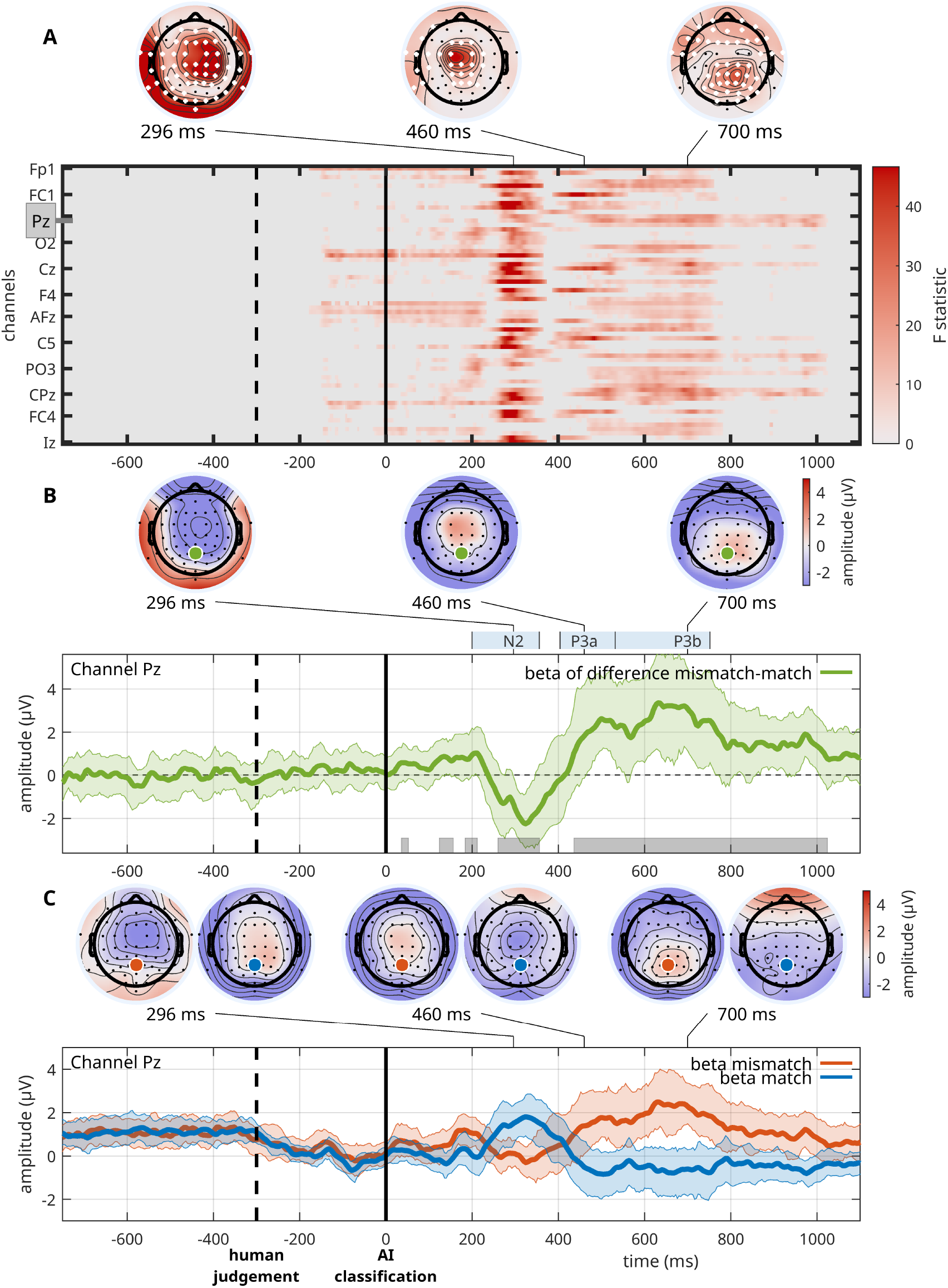
Mismatched vs. matched AI classification. **A**. ANOVA results comparing mismatch and match conditions, shown as a spatiotemporal heat map of significant F-values (red). Channels are sorted by proximity; Pz is marked and examined in panels B and C. Topographic maps are shown at 296, 460, and 700 ms after AI classification display. Electrodes in significant clusters are marked white; non-significant ones in black. Results are corrected using spatiotemporal clustering (*α* = 5%). **B**. Mismatch–match ERP time series (beta difference) at Pz, time-locked to AI classification (0 ms). Significant time points (from A) are marked in gray. Topographies at 296, 460, and 700 ms correspond to the N2, P3a, and P3b components. Curves show 20% trimmed means; shaded areas denote 95% Bayesian Highest Density Intervals (HDI). **C**. Beta time series at Pz for mismatch (red) and match (blue) conditions. Topographies for each condition are shown at the same three latencies, with Pz marked in red (mismatch) and blue (match). Data are shown as 20% trimmed means with 95% Bayesian HDIs.

Amplitude differences between mismatch and match conditions at the respective peaks were as follows: N2 = –1.34 *μ*V, 95% CI [–2.48, –0.09]; P3a = 1.83 *μ*V, 95% CI [–0.15, 4.19]; and P3b = 3.08 *μ*V, 95% CI [1.14, 4.94]. Figure 3C shows the ERP waveforms for match and mismatch conditions separately.

#### Processing AI classification depends on human judgment

We investigated whether neural responses to the AI classification (mismatch vs. match) were modulated by the participant’s preceding judgment of the image (fake vs. real), by testing the interaction term of the 2 × 2 repeated-measures ANOVA described above. The interaction contrast revealed a significant spatio-temporal cluster spanning all electrodes and time points from 192 to 888 ms (average F = 11.77, average p = 0.009; Figure 4A). To identify temporally specific effects within this window, we conducted post hoc t-tests on the interaction contrast using a more stringent cluster-forming threshold of *α* = 0.01 (supplementary Figures S4 to S6). This analysis revealed two significant time windows: one centered at 360 ms (within the N2 range) with a central-parietal topography, and a second at 484 ms (within the P3b range) with a broader distribution spanning frontal and occipital electrodes.

**Figure 4.**
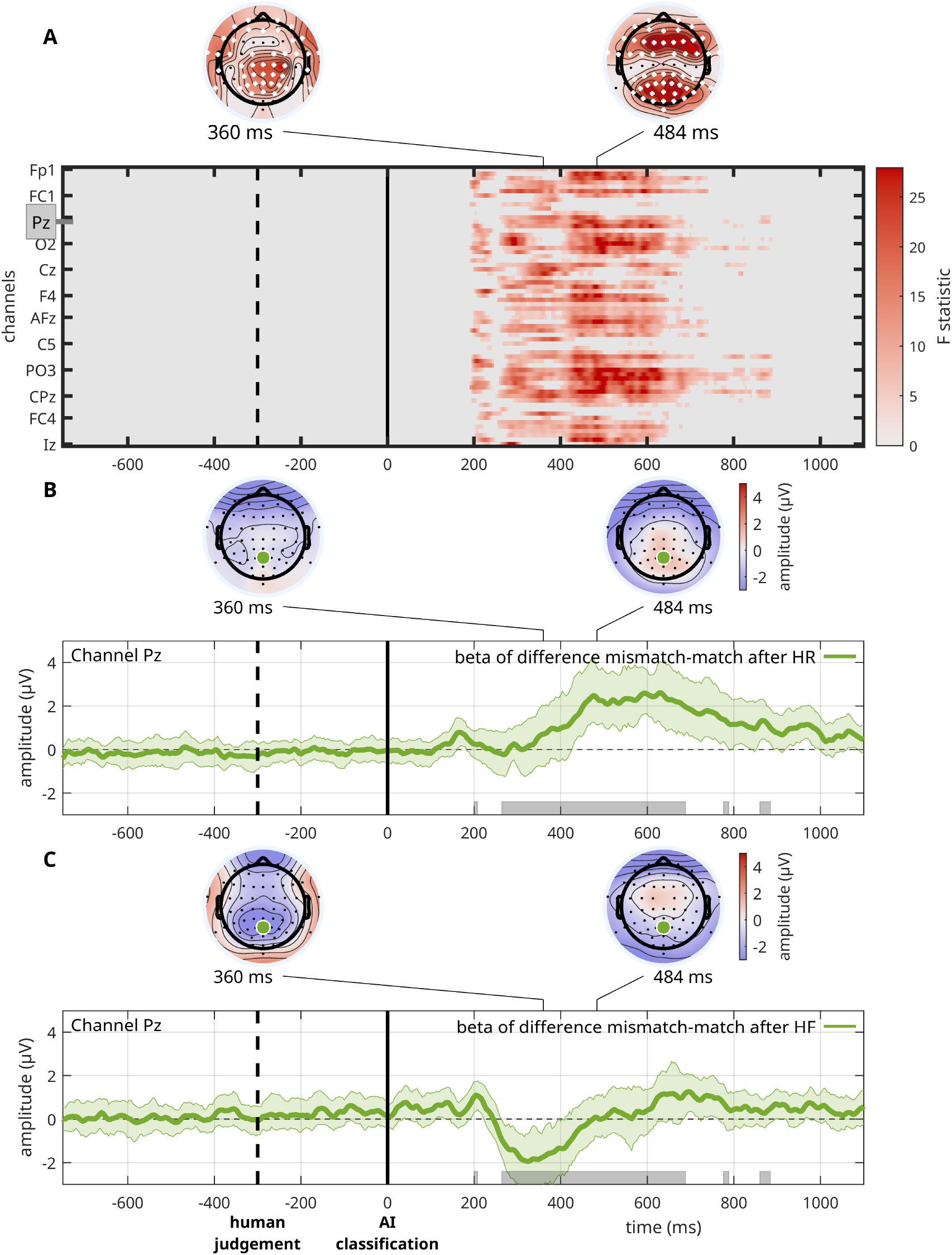
Interaction effects of AI classification (mismatch, match) and human judgment of images (fake, real). **A**. Interaction effect shown as heat map over channels and time points indicating significant F-values. Pz is marked for further description of the beta difference time courses in panels B and C. Topological distributions are presented at 360 and at 484 ms. Electrodes belonging to significant clusters are marked white; non-significant electrodes are marked black. **B**. Time series of beta mismatch-match difference wave when human judged an image as real at channel Pz with significant time points (as determined by the ANOVA results shown in panel A) marked in gray along the x-axis and topological distributions at 360 and 484 ms. Beta difference time series are presented as the 20% trimmed mean, shaded areas denote the 95% Bayesian Highest Density Interval. **C**. Time series of difference beta (mismatch-match) when human judged an image as fake at channel Pz, with significant time points marked in gray along the x-axis and topological distributions at same latencies. HF, Human Fake; HR, Human Real.

As illustrated by the condition-wise beta difference waveforms (Figure 4B and C), the mismatchminus-match ERP effect differed according to the participant’s prior judgment. Following a ‘fake’ judgment, the early negative deflection between 200–400 ms (N2 window) was larger, whereas the later positivity between 400–700 ms (P3b window) was minimal. Specifically, the mismatch-minus-match amplitude at 360 ms was –1.74 µV (95% CI [–2.99, –0.49]), and at 484 ms was 0.07 µV (95% CI [–0.80, 1.09]). In contrast, following a ‘real’ judgment, the early mismatch effect was attenuated (0.71 µV, 95% CI [–0.77, 2.04]), but a robust late positivity emerged at 484 ms (2.36 µV, 95% CI [1.13, 3.76]).

These findings indicate that the shape and timing of the mismatch–match ERP difference depend on the participant’s prior decision, with earlier effects more pronounced after ‘fake’ judgments and later effects stronger after ‘real’ judgments. This interaction suggests that the internal decision context modulates the temporal dynamics of how AI classification is being processed.

#### ERP responses of judging an image as fake or real

We tested whether the human judgment itself (fake vs. real) was reflected in the ERP responses by examining the second main factor of the 2 × 2 repeated-measures ANOVA. The contrast between ‘fake’ and ‘real’ judgments (i.e., HFAIF and HFAIR vs. HRAIR and HRAIF) revealed a significant spatio-temporal cluster spanning 34 electrodes from –728 to –412 ms relative to the onset of the AI classification (average F = 8.86, average p = 0.014; Figure 5A). Since the participant’s response was time-locked at –300 ms relative to AI onset, this cluster corresponds to –428 to –112 ms before the key press.

**Figure 5.**
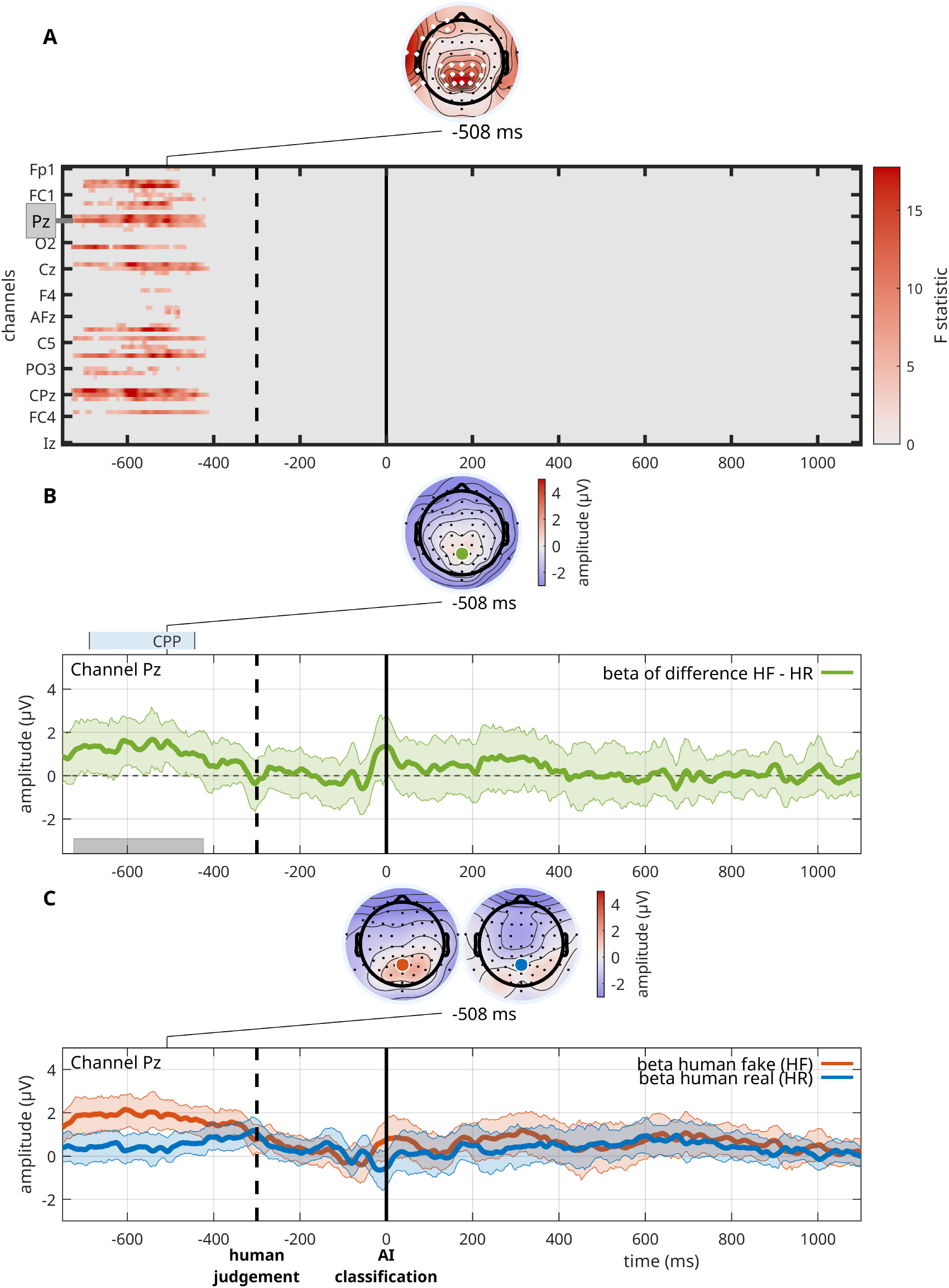
Human judgment of images (fake versus real). **A**. ANOVA statistical summary comparing human image judgment as fake or real shown as heat map indicating significant F-values in red over channels and time points. Pz is marked for further investigation of the beta time courses in panels B and C. Topological distributions are presented at −508 ms relative to AI classification. Electrodes belonging to significant clusters are marked white; non-significant electrodes are marked black; results are corrected with spatiotemporal clustering at an alpha level of 5%. **B**. Time series of difference beta for human judgment fake-real Pz, with time points belonging to the significant cluster marked in gray along the x-axis, and topological distribution at −508 ms. Beta time series are presented as the 20% trimmed mean, shaded areas denote the 95% Bayesian Highest Density Interval. **C**. Time series of betas at Pz for the human fake (red) and human real (blue) judgment conditions. Beta time series are presented as the 20% trimmed mean; shaded areas denote the 95% Bayesian Highest Density Interval. CPP, centro-parietal positivity; HF, Human Fake; HR, Human Real.

Post-hoc t-tests for the human judgment contrast, using a cluster-forming threshold of *α* = 0.01, identified a single significant cluster (supplementary Figure S7) and localized a centro-parietal positivity (CPP) peaking at –508 ms, with a topographical maximum over centro-parietal electrodes.

ERP waveforms at electrode Pz (Figure 5B and C) revealed that trials judged as ‘fake’ evoked higher amplitudes than those judged as ‘real’ during this image-viewing, pre-response window. At the peak time point of –508 ms, the fake–real amplitude difference was 1.6 *μ*V (95% CI [0.61, 2.49]).

### Neural processes of assessing and rating the AI’s reliability

We investigated whether changes in participants’ ratings of the AI’s reliability across the 20 blocks were reflected in the ERPs. To this end, we conducted another second-level analysis in LIMO using the same factorial design as in the ERP analysis reported above, but replaced the condition-based regressors with regressors that encoded block-wise changes in reliability ratings. These regressors estimated the covariance between ERP amplitude and participants’ evolving perceptions of AI reliability (see Methods for details).

Reliability ratings varied substantially across participants (Figure 6A to C; supplementary Figure S8). While many participants showed only minor fluctuations around their individual mean ratings, others exhibited pronounced upward or downward trends across blocks (Figure 6C). Distributions of block-wise changes were approximately normal for each participant, with Kullback–Leibler (KL) divergence values ranging from 0.03 to 0.1.

**Figure 6.**
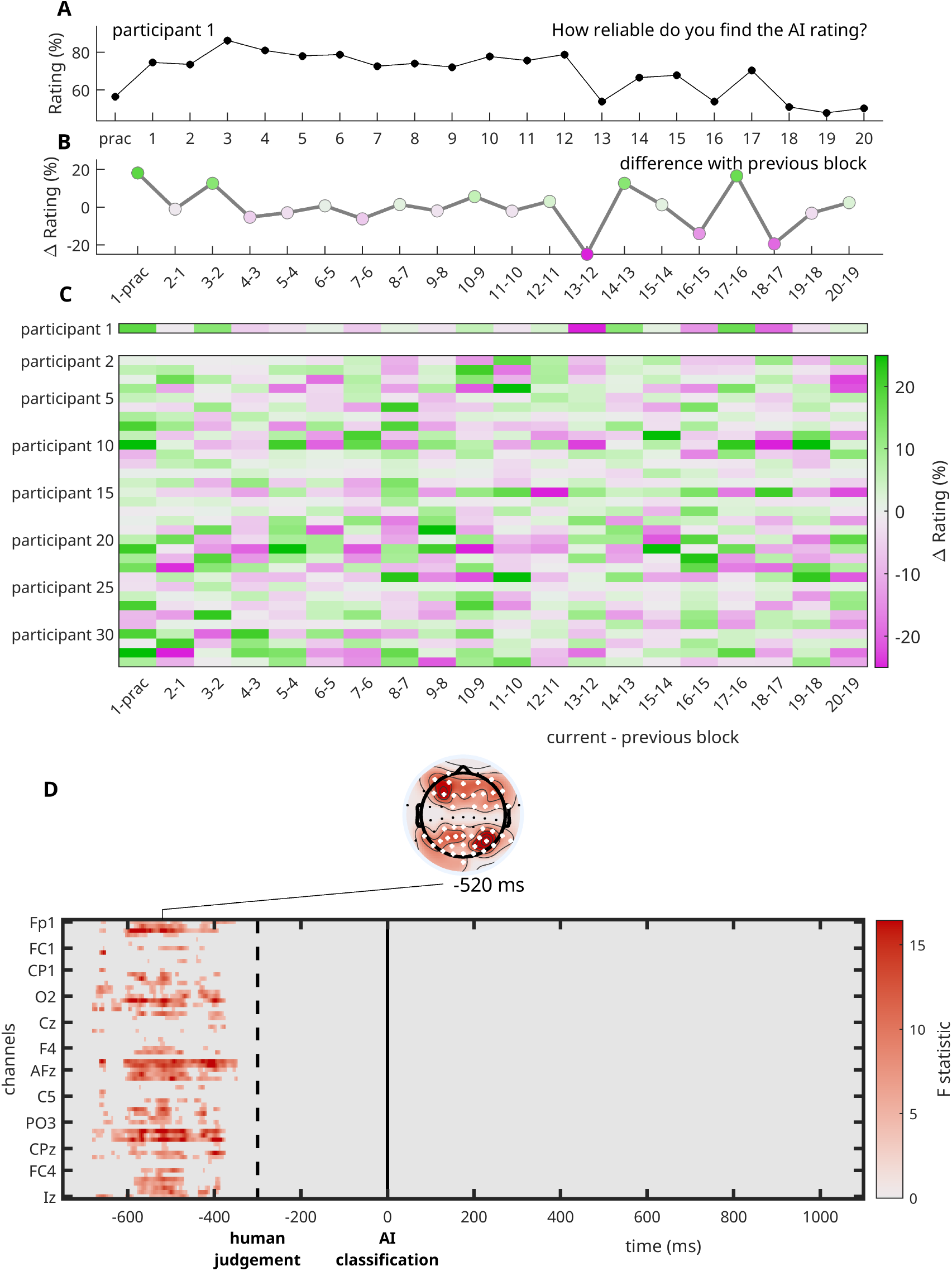
EEG amplitude co-varies with changes in perceived AI reliability. **A**. Reliability ratings (in %) across all blocks of experiment of participant 1 (one rating after each block, including the practice block). **B**. Difference in reliability ratings between blocks (e.g., reliability rating of block 1 - rating of practice block) for participant 1. Positive differences are colored green, negative differences purple. **C**. Overview of differences in reliability ratings between blocks for all participants. The first row depicts results of participant 1 and is identical to the data shown in panel (B) presented with a color map. The color map quantifies the difference in reliability ratings between blocks. **D**. ANOVA outcome for covariance between changes in reliability ratings and ERP amplitude over participants. Results are shown as heat map indicating significant F-values in red over channels and time points. Topological distributions are presented at −520 ms (before AI classification). In the topoplot electrodes belonging to significant clusters are marked white; non-significant electrodes are marked black; results are corrected with spatiotemporal clustering at an alpha level of 5%.

As shown in Figure 6D, changes in reliability ratings significantly influenced ERP amplitude. We identified one significant spatio-temporal cluster spanning 58 electrodes from –680 to –348 ms relative to the display of AI classification (average F = 8.33, average p = 0.016). The maximum F-value occurred at –520 ms, with a topographic distribution showing activity over both frontal and parietal electrodes. While this temporal window overlaps with the centro-parietal positivity (CPP) described in the preceding section, the topographic pattern here features a fronto-parietal split, which suggests a different temporally adjacent process.

To determine which conditions contributed most to this effect, we conducted post-hoc t-tests examining the condition-specific covariation between ERP amplitude and reliability rating changes (supplementary Figures S9 to S12). The strongest modulation was observed for trials in which participants judged the image as real and the AI classification was matching (HRAIR trials; supplementary Figure S9).

To illustrate the consistency and directionality of the effect across participants, we also computed trial-wise correlations between ERP amplitude (averaged per block) and changes in reliability ratings across blocks, for HRAIR trials at electrode Fz at –520 ms (supplementary Figure S13). While the main statistical inference was performed on beta estimates using cluster-based methods, these ‘raw’ correlations provide a complementary view of individual variability. Most participants (n = 25) showed a positive correlation, while 9 showed a negative correlation. This distribution illustrates a localized tendency towards a positive association between ERP amplitude and reliability updating at this specific time point and electrode.

### Human bias for fake or real image judgment influences processing of AI classification (mismatched, matched)

We examined whether individual differences in response bias influenced how participants processed AI classifications. Based on the response tendencies shown in Figure 1, we computed a continuous bias index ranging from –1 (consistent ‘real’ responder) to +1 (consistent ‘fake’ responder), reflecting each participant’s tendency to judge faces as fake.

We then performed a regression analysis between this response bias index and each participant’s beta estimates for three ERP contrasts: AI classification type (mismatch vs. match), human judgment (fake vs. real), and their interaction. This analysis tested whether response bias systematically influences ERP amplitude.

For the mismatch–match contrast, we identified four significant clusters where ERP amplitude covaried with response bias (Figure 7A). The first cluster spanned 14 electrodes from –96 to 292 ms (average F = 9.16, average p = 0.013), with a peak effect at 120 ms over frontal and parietal-occipital sites. The second cluster (464–784 ms, average F = 7.93, average p = 0.014) peaked at 548 ms over right frontal electrodes (AF4). The third cluster (492–784 ms, average F = 6.54, average p = 0.020) peaked at 636 ms over parietal electrodes (O1). The fourth cluster (808–884 ms, average F = 6.38, average p = 0.022) peaked at 836 ms over central-parietal regions (CP3). Post-hoc t-statistics revealed a consistent topographical pattern across the significant clusters, showing negative correlations at frontal sites and positive correlations at parietal sites (supplementary Figure S14). This spatial pattern complements the ANOVA results by illustrating the direction of the bias-related modulation across the scalp.

**Figure 7.**
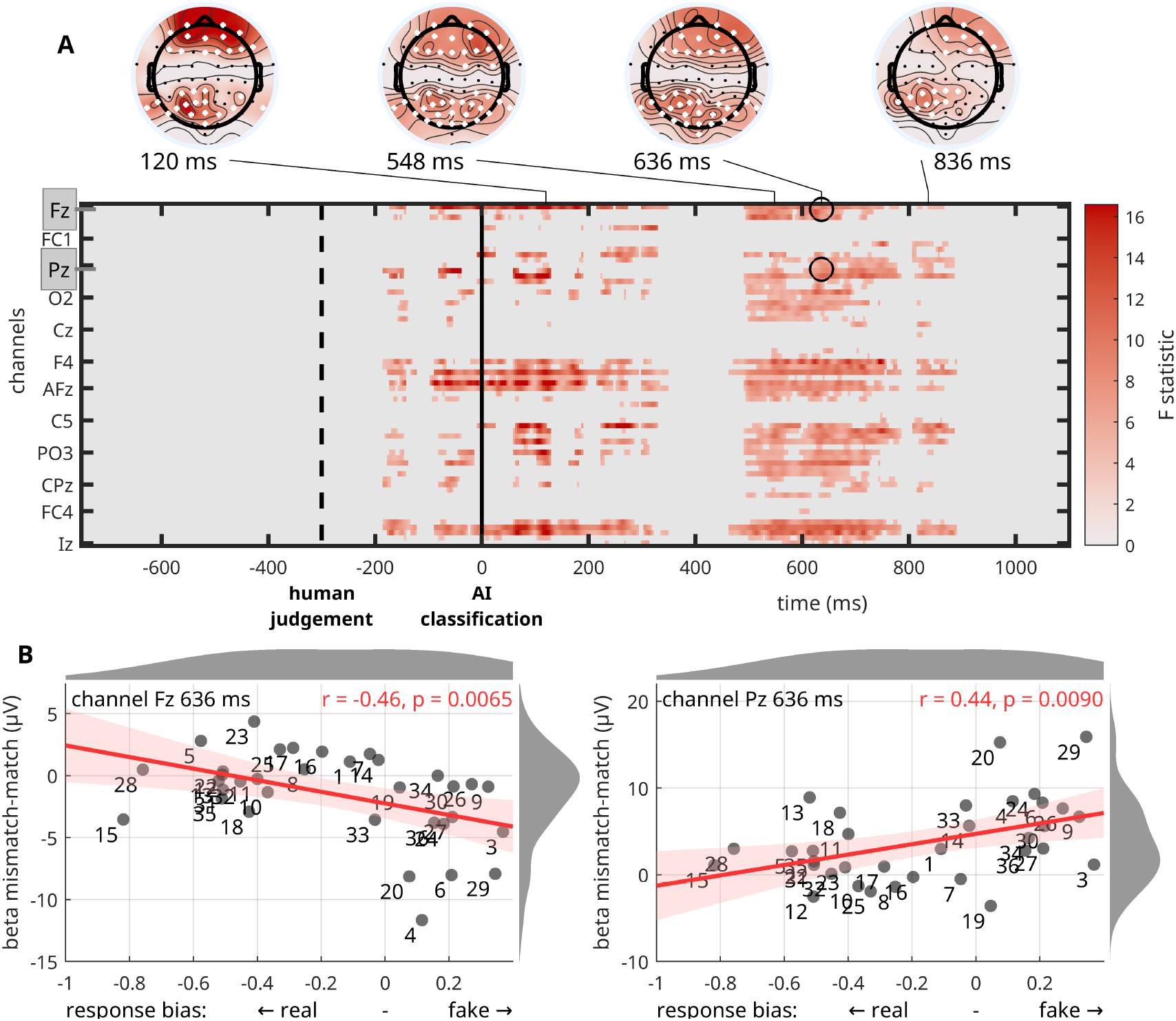
Regression results of human response bias for judging images as fake or real and mismatch-match ERP amplitude. **A**. F-statistic results from linear regression for the AI-classification-locked betas checking for correlations of human bias for judging images as real or fake and ERP (beta) amplitude of the mismatch-match difference wave. Results are shown as heat map indicating significant F-values in red over channels and time points. Channels Fz and Pz are highlighted for further investigation in panel B. Topological distributions are presented at four latencies (120 ms, 548 ms, 636 ms and 836 ms). Electrodes belonging to significant clusters are marked white; non-significant electrodes are marked black. Results are corrected with spatiotemporal clustering at an alpha level of 5%. Results are shown as heat map indicating significant F-values in red over channels and time points. Channels Fz and Pz are highlighted for further investigation in panel B. Topological distributions are presented at four latencies (120 ms, 548 ms, 636 ms and 836 ms). Electrodes belonging to significant clusters are marked white; non-significant electrodes are marked black. Results are corrected with spatiotemporal clustering at an alpha level of 5%. **B**. Correlations between bias for judging an image as fake and the amplitude of the difference mismatch-match beta at channel Fz (left) and Pz (right) at 636 ms. Gray dots represent participants, red line shows regression across participants, red shaded area the 95% CI. X-axis: values > 0 indicate a bias for fake, < 0 for real. Gray shaded areas at the edge of panels show the distribution of the response bias across participants (x-axis direction) and the distribution of the mismatch-match beta amplitude (y-axis direction).

This fronto-parietal split is illustrated in Figure 7B, showing ERP amplitudes at Fz and Pz at 636 ms. At frontal sites, participants with a stronger bias toward ‘fake’ judgments showed a greater negative deflection in the mismatch condition. At parietal sites, the same participants showed a greater positive deflection, suggesting that individual biases influenced the neural processing of AI classification outcomes in a topographically dissociable manner.

To determine whether these effects were driven more strongly by mismatch or match trials, we repeated the regression analysis separately for each condition (supplementary Figures S15 and S16). In the mismatch condition, bias effects were primarily observed in early time windows (–188 to 132 ms), whereas in the match condition significant clusters emerged later (from 308 to 1028 ms). This dissociation suggests that response bias may differentially influence the early detection of conflict and the later processing of agreement.

## Discussion

This study examined the neuro-cognitive dynamics of trust in a cooperative human–AI face classification task, in which the human first judged whether a face was fake or real, followed by an AI classification of the same image. We identified distinct neural signatures (ERPs) associated with human judgment, perception of the AI’s classification, and neural responses to agreement or disagreement between the two. These unfolded as a temporal sequence, beginning with internal decision formation and followed by external feedback processing. Disagreement from the AI (effectively a violation of expectation) elicited modulations of the N2, P3a, and P3b ERP components, reflecting conflict detection, attentional reorientation, and working memory updating. These ERP responses were not uniform but varied depending on the participant’s own judgment, indicating an interaction between external AI behavior and internal decision context. During the image viewing period and prior to the display of the AI classification, we observed a centro-parietal positivity (CPP), an ERP component associated with internal evidence accumulation and decision formation. This signal reflects the participant’s own decision-making process, independent of AI feedback. Additionally, we found that the amplitude of ERP responses co-varied with block-wise changes in perceived AI reliability in a similar image viewing, pre-AI-feedback time window. This finding suggests that internal decision dynamics may be consolidated in the updating of trust over time, even before external feedback is received. Finally, we found that individual differences in response bias modulated the neural processing of the AI classification, pointing to the role of personal heuristics in shaping how violations of expectation are interpreted. Together, these findings indicate that neuro-cognitive markers of trust in AI reflect a combination of internal evidence accumulation, feedback-related neural responses, and individual judgment biases.

### ERP signals reflect violations of internal expectations

Disagreement between the AI and the human’s judgment elicited a sequence of ERP components following the display of the AI’s classification. Compared to agreement trials, mismatch trials evoked a stronger fronto-central negativity around 300 ms, followed by a larger centro-parietal positivity between 400–800 ms (Figure 3). This pattern resembles the well-known N2–P3 complex typically observed in performance monitoring tasks, with comparable timing (200–750 ms) and topography (***De Visser et al***., 2018; ***Somon et al***., 2019).

The early negative deflection (N2) was not only modulated by mismatched AI classification but also by the participant’s own preceding judgment (Figure 4). This suggests that the brain’s initial response to disagreement depends on the interaction between external feedback and internal decision context. This N2 likely reflects a conflict detection or action–outcome violation signal, consistent with proposals that the medial frontal cortex generates prediction error signals when observed outcomes deviate from expected ones (***Alexander and Brown***, 2011; ***Ullsperger et al***., 2014).

The subsequent positivity from 400 ms onward most likely reflects a P3a–P3b complex. The P3a, with a more central scalp distribution, was sensitive to both mismatched AI classification and the participant’s preceding judgment, which implies reorientation of attention to unexpected or contextually salient feedback (***Polich***, 2007; ***Nieuwenhuis et al***., 2005). In contrast, the later parietal P3b was less influenced by the participant’s judgment and may index broader processes of outcome evaluation and working memory updating (***Donchin and Coles***, 1988; ***Polich***, 2012).

This ERP profile we found is consistent with prior work in human–automation interaction, where similar components have been interpreted as observational error-related negativity (oERN) and error positivity (oPe) in response to agent errors (***De Visser et al***., 2018; ***Somon et al***., 2019). Notably, the component latencies in our task were slightly delayed, likely due to increased perceptual and conceptual demands of judging face realism in the absence of ground-truth feedback.

These results align with the proposal posed in ***Ullsperger et al***. (2014), i.e. that the ERN, N2, and feedback-related negativity (FRN) constitute a family of temporally distinct error-monitoring signals, with the ERN typically response-locked, the N2 stimulus-locked, and the FRN feedback-locked. Despite these temporal differences, they may reflect a shared cognitive operation: detecting violations of internal expectations. In our case, a feedback-locked N2–P3 complex was observed, suggesting engagement of this broader action–outcome evaluation system in response to AI disagreement. The P3b, in particular, may reflect a slowly accumulating evaluative signal that supports outcome monitoring, context updating, or adaptive adjustment in uncertain environments (***Donchin and Coles***, 1988; ***Polich***, 2007; ***O’connell et al***., 2012).

Finally, given prior evidence linking the FRN/N2 to expectancy violation and surprise (***Hauser et al***., 2014; ***Amiez et al***., 2012; ***Jessup et al***., 2010), it is plausible that disagreement between the AI and the participant evoked a prediction error when the AI’s classification conflicted with the human’s internal judgment. This would support the interpretation that participants project a degree of subjective certainty onto the AI, and experience mismatch as a form of trust violation — even in the absence of objective correctness.

### Neural dynamics of human decision formation

During the image viewing period leading up to the human decision, we observed a slow, ramping signal over centro-parietal electrodes resembling the centro-parietal positivity (CPP; Figure 5), a well-established neural marker of evidence accumulation during decision formation ((***O’connell et al***., 2012; ***Kelly and O’Connell***, 2013; ***Loughnane et al***., 2016)). This component was stronger for trials in which participants judged the face as fake, suggesting that detecting artificial faces demanded more accumulated evidence than recognizing real ones. The topography and temporal profile of the signal are consistent with internal commitment dynamics (evidence building toward a threshold) prior to response ((***Shadlen and Kiani***, 2013)). These findings support the idea that the CPP may serve as a readout of participants’ internal decision formation, independent of any external feedback or subsequent AI response.

### Neural markers of trust evolution

Surprisingly, we also found during the image viewing period that ERP amplitude was modulated by block-wise changes in perceived AI reliability (our behavioral proxy for trust). The time period of this effect was unexpected because it emerged prior to the display of the AI classification, during the image viewing interval. This time window is typically associated with internal decision formation (i.e., the CPP, as discussed above). The effect was evident at both frontal and parietal sites and was most pronounced in trials when the human judgment was real and the AI classification matched (HR-AIR).

The topography suggests a distributed network involving both evidence accumulation and metacognitive monitoring ((***Fleming and Dolan***, 2014; ***Bang and Fleming***, 2018)), possibly reflecting participants’ integration of internal certainty with evolving beliefs about the AI’s reliability. That is, although participants were explicitly asked to evaluate the AI, the data indicate that they may have drawn (perhaps subconsciously) on their own sense of decision confidence. These findings suggest that participants maintained an internal ‘record’ of their decisions, accessible at a meta-cognitive level, to be used later to guide updates to their impression of the AI over time.

### Individual decision bias modulates neural responses to AI classification

Participants differed in their tendency to judge faces as fake or real. This response bias correlated with the mismatch-minus-match ERP difference (Figure 7, supplementary Figure S14), with a striking topographical split: negative correlations at frontal sites and positive correlations at parietal sites. Two temporal clusters emerged, an early effect from 96 to 292 ms, and a later effect from 464 to 884 ms relative to AI classification display.

The early effect, primarily driven by mismatch trials (supplementary Figure S15), extended into the latency range of the N2 and was characterized by increased frontal negativity in participants with a stronger bias toward ‘fake’. The later effect, mainly driven by match trials (supplementary Figure S16), overlapped with the P3b and showed enhanced parietal positivity in those same participants. This dissociation suggests that bias shapes distinct cognitive stages of AI classification processing: early conflict monitoring when the AI disagrees, and later belief confirmation when the AI aligns with expectation.

These findings align with predictive coding accounts, in which internal models shape how feedback is evaluated. Participants with strong priors (e.g., that most faces are fake) may process AI classifications differently depending on whether they violate or confirm those expectations. The variability in the strength and topography of these effects across individuals (Figure 7B) reflects heterogeneity in how bias influences evaluative processing and outcome monitoring.

These findings are consistent with the predictive coding framework, which proposes that internal models shape the neural evaluation of feedback (***Friston***, 2010; ***Gilboa and Marlatte***, 2017; ***Alexander and Brown***, 2011). Participants with a strong bias toward ‘fake’ or ‘real’ classifications may process AI feedback differently depending on whether it violates or confirms their expectations, which influences both early conflict-related responses and later outcome evaluation. While our data do not directly demonstrate predictive coding mechanisms, they do not exclude such interpretations and point toward a role of internal priors in shaping ERP dynamics. The observed inter-subject variability (Figure 7B) suggests heterogeneity in how these biases affect evaluative strategies, which could be a promising direction for future research.

### A layered framework for trust formation: evaluative, decisional, and meta-cognitive dynamics

Taken together, our findings reveal that trust in human–AI interaction is not a singular process in the brain, but rather emerges from the interplay of multiple neural processes operating at different temporal and cognitive levels. Rapid event-related potentials (N2, P3a, P3b) reflect the immediate evaluation of AI feedback, including detection of agreement or conflict. Concurrently, a decision-related signal (the CPP) indexes the internal accumulation of evidence leading up to the human’s judgment. Crucially, ERP amplitude during this same period co-varied with subsequent changes in perceived AI reliability, suggesting that trust updates draw not only on external feedback, but also on internal assessments of belief certainty. Finally, the influence of individual decision biases on feedback-locked ERP components highlights the potential role of internal priors in shaping how agreement and disagreement are processed. Together, these results support a layered framework in which trust is continuously formed, modulated, and re-evaluated through the coordination of evaluative, meta-cognitive, and decisional systems.

In addition to these classical interpretations, we note one speculative perspective proposed in prior theoretical work on Quantum Cognition theory (***Busemeyer and Bruza***, 2025; ***Pothos and Busemeyer***, 2013; ***Bruza et al***., 2015). This account proposes that the CPP response may reflect an indeterminacy in the participant’s cognitive state, consistent with principles of Quantum Cognition. That is, participants may not be in a definite cognitive state at all times (e.g., judging the image as either fake or real), but rather in an indeterminate superposition of both possibilities. In other words, prior to making an explicit judgment, the cognitive state may simultaneously represent both ‘fake’ and ‘real’ outcomes. Once a decision is made and reported, this superposed state collapses into a definite cognitive state corresponding to a single decision level. Thus, prior to being measured, both levels of the image judgment coexist as propensities in the cognitive state, and this cognitive indeterminacy may be reflected in the ERP signal we observed. Similarly, this principle of indeterminacy may also extend to the meta-cognitive processes involved in trust updating.

## Limitations

This study has several limitations. While our sample size is comparable to prior EEG research, a larger cohort would enable more fine-grained analyses of inter-individual variability—particularly in distinguishing participants with a general tendency to favor or distrust the AI (“AI proponents” vs. “AI skeptics”). Additionally, incorporating complementary physiological measures such as pupil diameter or eye-tracking could provide further insight into attentional engagement, arousal, or strategic shifts during decision-making.

In the present design, reliability ratings were collected only at the end of each block. Capturing trial-wise reliability ratings could help clarify whether the observed ERP–trust correlations reflect explicit meta-cognitive awareness or instead index implicit confidence dynamics. However, such a design would significantly increase the duration of the task and may introduce additional confounding factors. Furthermore, including trial- or block-wise ratings of participants’ confidence in their own judgments (“confidence-in-self”) would allow a clearer dissociation between trust in the AI and trust in oneself, two related but distinct constructs likely to influence neural dynamics in different ways.

## Outlook

Our findings open several promising directions for future research. Computational frameworks such as the Hierarchical Gaussian Filter (HGF) (***Mathys et al***., 2014) provide principled models of how internal certainty and external feedback jointly shape belief updates over time. In this context, EEG signals, particularly components like the CPP and feedback-locked potentials, could be used not merely as outcome measures but as real-time inputs to constrain or update latent belief states. This opens the door to closed-loop systems in which neural markers dynamically inform adaptive AI behavior, potentially enabling more robust and personalized trust calibration.

Quantum cognitive models offer a complementary formalism for capturing indeterminate belief states that evolve dynamically prior to resolution (***Pothos and Busemeyer***, 2013; ***Busemeyer and Bruza***, 2012; ***Bruza et al***., 2015; ***Busemeyer and Bruza***, 2025). These models may be especially valuable when trust must be inferred in the absence of explicit feedback, or when internal evaluations unfold in context-dependent, nonlinear ways.

Such frameworks align with modern meta-cognitive theories, where internal signals of certainty, conflict, or expectation violations continuously shape how beliefs are formed and revised (***Fleming and Daw***, 2017). Future studies could incorporate trial-level adaptive paradigms (***Nassar et al***., 2010) to investigate how trust evolves under dynamically shifting conditions; this represents an essential step toward real-world applications in critical human–AI teaming scenarios such as defense, autonomous driving, or clinical decision support systems. In these settings, EEG-derived markers could act as continuous, individualized indicators of belief resolution, potentially enabling intelligent systems to adapt in sync with users’ evolving internal states. This vision is supported by EEG–fMRI evidence linking confidence tracking to the ventromedial prefrontal cortex (vmPFC), a key region implicated in adaptive control and belief integration (***Lebreton et al***., 2015).

## Conclusion

In summary, this study shows that neural markers of decision making, feedback processing, and individual bias each contribute to how humans evaluate the reliability of an AI partner. Trust was not merely reactive to the AI’s outputs, but emerged from an interaction between internal decision signals, prior expectations, and perceived reliability of the AI. This research contributes to a growing understanding of the neurophysiological basis of trust dynamics in human–AI teams.

## Methods and Materials

### Participants

Healthy adults between 18 and 34 years of age participated in this study (in total 36 participants, 50% female, mean age 26 years). All participants gave written informed consent prior to participation and they received financial compensation in the form of a non-specific gift voucher (AUD 25). Participants were right-handed, had normal or corrected-to-normal vision, had no musculoskeletal injuries of the upper limbs and back, had no neurological or psychiatric disorders and were not under any neurotropic medication. As described above, data from 2 participants were excluded (one for non-completion, and one for non-compliance) and data from 34 participants were used for analyses. Results of one participant were previously described in ***Roeder et al. (2023***). The experimental protocol was approved by the Human Research Ethics Committee of Queensland University of Technology (LR 2022-5210-10103) and by the U.S. Air Force Human Research Protection Office (FWR20220251X) in accordance with the Declaration of Helsinki. Throughout this work, we followed guidelines for reproducible research as outlined by the OHBM COBIDAS MEEG committee (***Pernet et al***., 2020).

### Experimental protocol

Participants took part in an image classification task. They had to judge whether images of human faces are ‘fake’ (AI-synthesized) or ‘real’, and were afterwards (300 ms) presented with the classification of a simulated AI agent of that same image. After each block of such 28 trials, they rated the reliability of the simulated AI agent’s image classification. (For simplicity, we refer to the simulated AI agent as AI in this article.) Participants were instructed before and during the experiment that they were interacting with an AI agent. Only after the experiment, were they debriefed to clarify that the ‘AI agent’ was ‘simulated’, that is, a pre-programmed experiment. The experiment was completed in one session of approximately two hours which included briefing and preparation of the EEG. Figure 8 gives an overview of the experimental design.

**Figure 8.**
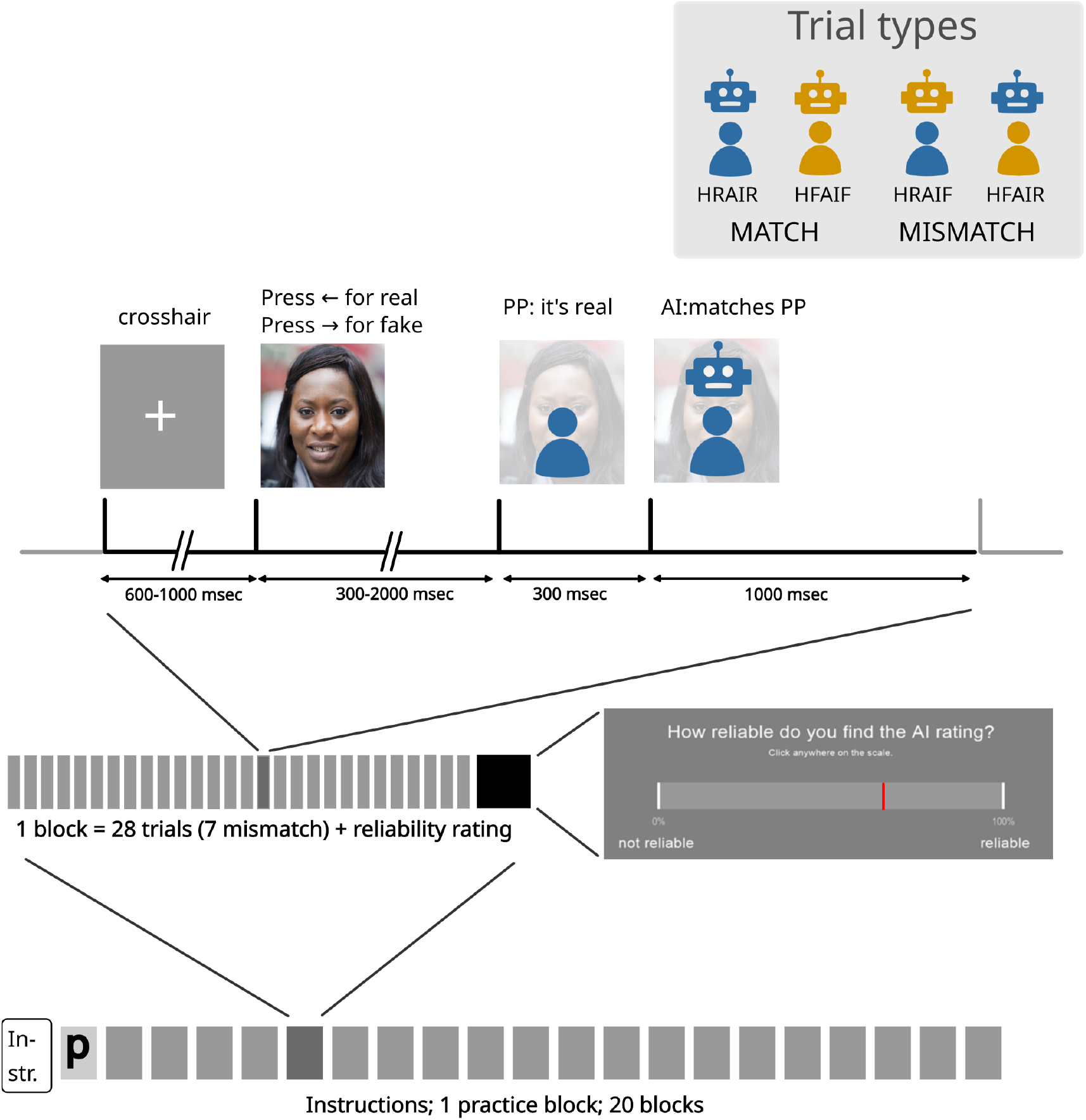
Overview of the experimental design. Trial structure (top), block structure (middle), experimental procedure (bottom). The trial structure represents one example when participants judged an image as real and the AI classification matched their judgment; see text for details on fake and mismatch conditions. The trial types are depicted top-right and are as followd: HRAIR, Human-Real & AI-Real; HFAIF, Human-Fake & AI-Fake; HRAIF, Human-Real & AI-Fake; HFAIR, Human-Fake & AI-Real.

Following instruction screens and a practice block of 28 trials, the experiment consisted of 20 blocks of 28 trials each (560 trials in total). Each trial started with a fixation cross in the centre of the screen lasting between 0.6 to 1 s. Subsequently, the image stimulus was displayed until the participant pressed a response key (judgment of real or fake) or for a maximum of 2 s. Upon pressing a response key, their own judgment was displayed on the screen: either a blue human icon if they judged the image as real, or a yellow human icon if they judged the image as fake. 300 ms after the human icon display, the AI classification was also displayed on the screen: either a blue robot icon if classified as real, or a yellow robot icon if classified as fake. Both the human and robot icon remained on the screen for 1 s before the next trial began or the reliability slider at the end of each block was displayed.

After each block (as well as before and after the practice block), participants were asked to rate the reliability of the AI classifications on a continuous scale with a slider from 0% to 100% (Figure 8) by using the mouse. Completing the reliability rating was self-paced. A response was logged by clicking the mouse button, but participants could not log a response without first moving the mouse in order to prevent lack of responding. After participants logged their response it was displayed on the screen for 2 s; participants could not change their response after it was first logged. After completion of the reliability slider, participants were invited to a short self-paced break. A longer self-paced break was offered after blocks 7 and 14.

The images used for the current study were drawn from the data set in (***Nightingale and Farid***, 2022). We selected 294 (50%) AI-synthesized (fake) faces and 294 (50%) real faces from their data set ensuring diversity across gender, age and ethnicity, and according to those most accurately identified as fake or real as rated in (***Nightingale and Farid***, 2022). Within each block of 28 trials, half of the images (14) were real and half (14) were fake. The order of images was randomized across trials, blocks and participants, and each image was displayed once only (no repeats) to each participant. Participants classified the images using the left and right arrow keys on a standard keyboard. Half of the participants had the left key assigned for the real rating and the right key for fake, and vice versa for the other half (counterbalanced order).

The experiment contained two conditions: match and mismatch. In the match condition, the AI classification matches the participant’s, i.e. if the participant judges an image as real, the AI agent also classifies the image as real. Vice versa, if the participant judges an image as fake, the AI also classifies it as fake. In the mismatch condition, the AI classification is always different from the participant’s classification. That is, if a participant judges an image as real, the AI classification is fake (and vice versa if a participant chooses fake). Within each block, 25% of trials were the mismatch condition and 75% were the match condition, and the order of match and mismatch trials was randomized in each block.

The experiment was run on a 64-bit Microsoft Windows 10 PC. The images and classification icons were displayed against a grey background using the PsychoPy3 standalone software version 2022.1.4 (***Peirce et al***., 2019) onto a monitor (with a 1920 × 1080 pixels resolution and a 60 Hz refresh rate) located 60 cm away from the participant, and mirrored onto a monitor for the experimenter. They consisted of images (6×6 cm, at a 5.7° visual angle) and overlaid feedback-icons (human icon displayed in the bottom half of the image, robot icon displayed in the top half of the image). Figure 8 shows an example of how the screen looked. Participants were seated in a chair that can be adjusted in height so that their eye height matched the centre of the monitor. The monitor could also be adjusted in height. The experiment was conducted in a shielded room. The experimenter was in the room with the participant, separated by a few meters and an opaque curtain.

### Data acquisition

Electroencephalographic (EEG) data were recorded using a 64-channel actiCAP system with active electrodes (BrainProducts GmbH, Gilching, Germany) placed according to the extended international 10–20 system (***Oostenveld and Praamstra***, 2001), with FCz as the recording reference and Fpz as the ground. The cap was positioned by centering the Cz electrode on the axes nasion to inion and left and right preauricular points. EEG signals were recorded continuously using the BrainProducts LiveAmp amplifier and BrainVision Recorder software (version 1.25.0101), with a 500 Hz sampling frequency on an independent recording PC running Windows 10. The triggers from the PsychoPy PC were sent to the EEG system via USB port that mirrors a parallel port and the BrainProducts TriggerBox and Sensor and Trigger Extension. At the beginning of the session, the experimenter fitted the participant with the EEG cap. Recording sites on the scalp were abraded using the Nuprep paste. EEG electrodes were attached onto the cap, filled with conductive gel (Supervisc, BrainProducts GmbH, Germany), and adjusted until impedances were below or close to 10 kOhm. Before and during the experiment, the participant was instructed to relax shoulders, jaw, and forehead and to minimise swallowing.

### Data analysis

#### Behavioral data

For all behavioral data analysis, we only used data from trials which were kept in the final ERP analysis after pre-processing and cleaning of EEG recordings.

#### Accuracy of image judgments

We counted the total number of valid image judgments (key presses) per participant, as well as the number of images judged as fake and the number of images judged as real. This way, we were able to calculate each participant’s accuracy in judging the images, and also their response bias (i.e., preference for judging the face as fake or real). Each participant had their own bias value between −1 and 1, defined as:

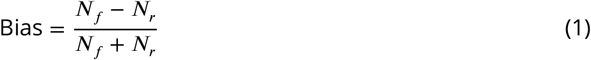

where:

- *N*_*f*_ is the number of responses where the participant judged the face as *fake*,
- *N*_*r*_ is the number of responses where the participant judged the face as *real*.

The bias index ranges from –1 to 1, with positive values indicating a preference toward fake responses, negative values indicating a preference toward real responses, and 0 indicating no response bias.

#### Response times

We analyzed the response time per participant by means of the time that a participant took to press the relevant fake or real key after an image was first presented. For this, we used the LiveAmp triggers in the ERP data to calculate response times and validated this with the response times logged by PsychoPy. To assess the effects of judgment type and individual response bias on response time (RT), we checked mean response times within participants (fake vs real) with a paired t-test. To assess whether this difference was interacting with response bias, we conducted a 2 (judgment: fake, real; within-subjects) × 2 (bias: fake-biased, real-biased; between-subjects) mixed ANOVA; for each participant, mean RTs were calculated separately for fake and real judgments. Participants were grouped into bias categories based on their overall tendency to respond fake or real across trials.

#### Reliability ratings

To track reliability ratings, we calculated changes in reliability ratings between blocks for each participant. To this end, we subtracted scores between adjacent blocks (e.g., rating block 2 - block 1). Resulting positive values indicate an increase in reliability perception, and negative values indicate a decrease in reliability perception. We used these changes in reliability ratings for further analyses with the ERP data.

#### EEG pre-processing

EEG data were pre-processed in MATLAB (version 2019b) and EEGLAB (version 2022.1) (***Delorme and Makeig***, 2004). To preserve the integrity of the underlying neural signal, we used an approach inspired by the PREP pipeline strategy (***Bigdely-Shamlo et al***., 2015), that uses component-based subtraction rather than signal replacement. This approach leaves lower and higher frequencies untouched and works in the continuous time domain; we consider epoching and filtering part of the analysis performed separately. The following formula indicates how the signals were preprocessed

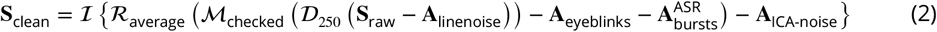

**S**_raw_ Raw EEG signal.

**A**_linenoise_ Estimate of line noise artifacts at 50, 100, and 150 Hz.

**𝒟**_250_ Downsampling to 250 Hz.

**ℳ**_checked_ Automated time segment and whole-channel rejection with visual inspection for confirmation.

**A**_eyeblinks_ Estimate of artifacts related to eyeblinks (from ICA eyenlink components).

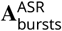 High-amplitude transient artifacts removed using Artifact Subspace Reconstruction.

**ℛ**_average_ Re-referencing to average reference.

**A**_ICA-noise_ Residual non-neural noise components identified via Independent Component Analysis.

**S**_clean_ Cleaned EEG signal.

**ℐ**Interpolation of previously removed channels using spherical interpolation.

The raw EEG signal, denoted **S**_raw_, was first cleaned of line noise artifacts (**A**_linenoise_), including peaks at 50, 100, and 150 Hz. The result was downsampled to 250 Hz using 𝒟_250_ to reduce data dimensionality. An automated preprocessing step, ℳ_checked_, was then applied using the clean_rawdata plugin in EEGLAB, which identified and removed unsalvageable channels and time periods based on statistical criteria (e.g., flatlining, low correlation, high variance, or extreme amplitudes). All rejections were visually reviewed and confirmed.

Eyeblink-related signal components (**A**_eyeblinks_) were estimated using the infomax ICA algorithm with PCA option, and ICLabel v1.6 (the threshold for the eye category was set to 0.9, i.e. with at least 90% confidence) (***Pion-Tonachini et al***., 2019). All other types of components (artifactual and brain) were retained in the data at this stage. Transient high-amplitude burst components 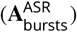 were estimated in the time domain using Artifact Subspace Reconstruction (ASR), also implemented in clean_rawdata, and subtracted from the data. The threshold for burst removal was set to 10 (standard deviations of PCA-extracted components). Data were then re-referenced to the average reference (ℛ_average_), and residual non-neural components identified via ICA (**A**_ICA-noise_) were likewise projected out. Channels removed earlier were interpolated using spherical splines. The resulting signal **S**_clean_ was used for all subsequent analyses.

Prior to performing the ICA steps to estimate the eyeblink and artefactual signal contributions, a 1 Hz high-pass filter was used. At each step of the analysis, the artefactual signal or component were estimated and saved to disk. Finally, the clean signal **S**_clean_ was calculated according to the subtraction in Equation 2, by loading the different components and subtracting them from the raw signal. This resulted in a clean, but unfiltered signal **S**_clean_.

#### EEG event-related analysis

After the EEG signals were cleaned, data were band-pass filtered between 0.3 Hz and 45 Hz, and segmented around the display of the AI classification icon on screen (Figure 8). The trials ranged from −750 ms before to 1100 ms after relative to this event. This is to identify ERPs time-locked to this event and to also explore the ERPs before the human image judgment (key press). Trials in which participants did not press the response key in time (>2 s), as well as trials in which participants pressed the response key in <0.35 s were removed from analyses. Overall, less than 1% of trials were excluded as almost all of the response times were within this timeframe. Baseline correction was applied based on a period between −500 ms to 0 ms relative to the presentation of the face (image stimulus), i.e. during the time when the fixation cross was displayed. We chose this baseline because the mental load is the same for all trials during this period (if we had chosen a baseline based on the response, this would potentially average out processes related to evaluation of the realness of the presented face).

Analysis of average ERPs over trials was performed using hierarchical linear model approach, as implemented the LIMO EEG toolbox (v3.3) (***Pernet et al***., 2011). The LIMO toolbox fits a regression model based on each participant’s trial-level data, at each time point and electrode (i.e., estimated beta parameters quantifying the condition-specific partial contribution to the full EEG signal) using a weighted least-squares approach that penalizes outliers. The estimated beta parameters are subsequently used in an appropriate second-level (group) statistical model to assess condition effects at group level.

At the first (participant) level, a general linear model (GLM) was set up with nine regressors, encoding the following conditions: 1. Human-Real & AI-Real (HRAIR), 2. Human-Fake & AI-Fake (HFAIF), 3. Human-Real & AI-Fake (HRAIF), 4. Human-Fake & AI-Real (HFAIR). We were also interested in assessing whether variation in the EEG co-varies with changes in reported trust levels (reliability ratings) across these conditions. We therefore seeded the model with additional continuous regressors that encode for these differences, creating the following additional regressors: 5. change in reliability ratings between blocks for HRAIR trials 6. change in reliability ratings between blocks for HRAIF trials, 7. change in reliability ratings between blocks for HFAIF trials, 8. change in reliability ratings between blocks for HFAIR trials. The last regressor (9.) encoded the baseline ERP, i.e. the average waveform common across all trials. Beta parameter estimates were obtained using trial-based Weighted Least Squares (WLS) (***Pernet et al***., 2022).

At the second level (group), condition effects were examined at each channel and time point using three types of analyses. First, we investigated differences in ERP amplitude between conditions with a 2×2 repeated-measures ANOVA with two factors: 1. AI classification (mismatched, matched), 2. human judgment (fake, real). The ANOVA also permits to test for interactions between these two factors. This model was seeded with the first four beta parameter estimates from the first (participant) level GLM (HRAIR, HFAIF, HRAIF, HFAIR). Post-hoc t-tests were performed with second-level one-sample t-tests seeded with contrast estimates calculated at first level: [−1, −1, 1, 1] for human judgment (yielding the ERP response estimate for fake vs real), [1, −1, 1, −1] for AI classification (yielding the ERP response estimate for mismatched vs matched), and [−1, 1, 1, −1] for the interaction of AI classification and human judgment. Results from post-hoc t-tests are reported in the supplementary material. Second, we investigated the co-variation between ERP amplitude and changes in reliability ratings. To this end, we set up another 2 × 2 repeated-measures ANOVA with factors 1. human judgment (fake, real), 2. AI classification (mismatched, matched), 3. interaction of the two main effects. In this model, we used the fourth to eighth beta parameter estimates from the first-level GLM (change in reliability ratings for HRAIR, change in reliability ratings for HFAIF, change in reliability ratings for HRAIF, change in reliability ratings for HFAIR). Since we are only interested in co-variance of the ERP amplitude, we created a F-constrast spanning all conditions as a 4 × 4 matrix with ones on the diagonal to check for covariance across conditions. To perform post-hoc t-tests on each condition separately, second-level (group) t-tests were calculated seeded with contrast estimates calculated at first level spanning a single condition (for example, [1, 0, 0, 0] for HRAIR). The third group analysis investigated whether the response bias of the participants co-varied with ERP amplitude of the mismatch-match difference wave. For this we constructed a regression model seeded with the contrast parameters estimated at first level encoding for mismatch-match ERP amplitude [−1, −1, 1, 1], and with the participant-wise regressor for response bias defined by Equation 1. We calculated post-hoc t-tests investigating the correlation of response bias within mismatch-only trials and match-only trials.

#### Statistical inference

All statistical tests (second-level repeated measures ANOVA, t-tests, and regressions) were performed with bootstrapping, to check for significant effects in a 2D matrix spanning channels (64 total) and time (relative to the AI classification). The type I error rate (due to multiple comparisons) was controlled via spatiotemporal cluster-based inference at an alpha level of 5%, according to guidelines outlined in the LIMO toolbox (***Pernet et al***., 2015). For each significant cluster, we report the time span and the channels enclosed within that cluster. Although we cannot make inferences about specific points in channel-time space, we report the time point at which the average F-value (or t-value) across all significant electrodes reaches its maximum for descriptive purposes. This provides an approximate indication of the cluster’s temporal peak but should not be interpreted as a separately significant time point.

Effect sizes (comparable to grand average difference ERPs) were calculated via contrasts of beta parameter estimates. To reduce the influence of outliers and skew, we used a 20% trimmed mean of the weighted beta estimates across participants, as implemented in LIMO EEG’s robust group analysis pipeline (***Pernet et al***., 2011). This approach improves robustness to non-normality and provides more reliable estimates of central tendency under EEG noise conditions. To quantify uncertainty, Bayesian 95% credible intervals were estimated using the posterior distribution over group-level parameters, providing an interpretable range within which the true effect is likely to lie (***Pernet et al***., 2011).

## Data Availability Statement

The raw EEG datasets contain identifiable participant information and cannot be shared publicly due to ethical and privacy constraints. The data together with the complete MATLAB analysis pipeline and code, are available from the corresponding authors upon reasonable request. The behavioral data has been used in our previous work on behavioral modeling in ***van der Meer et al. (2025***) and is available on github: https://github.com/qi-research-group/QRW_Markov_Walk.git.

## Acknowledgments

This material is based upon work supported by the Air Force Office of Scientific Research under award numbers: FA9550-22-1-0005, FA9550-23-1-0258. We thank Dr Michael Breakspear and Dr Saurabh Sonkusare for their thoughtful feedback on the manuscript and Dr Pat Johnston for insightful discussions during the initial design of the study.

## References

Alexander WH, Brown JW. Medial prefrontal cortex as an action-outcome predictor. Nature neuroscience. 2011; 14(10):1338–1344.

Amiez C, Sallet J, Procyk E, Petrides M. Modulation of feedback related activity in the rostral anterior cingulate cortex during trial and error exploration. Neuroimage. 2012; 63(3):1078–1090.

Bang D, Fleming SM. Distinct encoding of decision confidence in human medial prefrontal cortex. Proceedings of the National Academy of Sciences. 2018; 115(23):6082–6087.

Bigdely-Shamlo N, Mullen T, Kothe C, Su KM, Robbins KA. The PREP pipeline: standardized preprocessing for large-scale EEG analysis. Frontiers in neuroinformatics. 2015; 9:16.

de Bruijn ER, von Rhein DT. Is your error my concern? An event-related potential study on own and observed error detection in cooperation and competition. Frontiers in neuroscience. 2012; 6:8.

Bruza PD, Wang Z, Busemeyer JR. Quantum cognition: a new theoretical approach to psychology. Trends in cognitive sciences. 2015; 19(7):383–393.

Busemeyer JR, Bruza P. Quantum Models of Cognition and Decision: Principles and Applications. No. 2nd, Cambridge University Press; 2025.

Busemeyer JR, Bruza PD. Quantum models of cognition and decision. Cambridge University Press; 2012.

Carp J, Halenar MJ, Quandt LC, Sklar A, Compton RJ. Perceived similarity and neural mirroring: evidence from vicarious error processing. Social neuroscience. 2009; 4(1):85–96.

De Visser EJ, Beatty PJ, Estepp JR, Kohn S, Abubshait A, Fedota JR, McDonald CG. Learning from the slips of others: Neural correlates of trust in automated agents. Frontiers in human neuroscience. 2018; 12:309.

Delorme A, Makeig S. EEGLAB: an open source toolbox for analysis of single-trial EEG dynamics including independent component analysis. Journal of neuroscience methods. 2004; 134(1):9–21.

Donchin E, Coles MG. Is the P300 component a manifestation of context updating? Behavioral and brain sciences. 1988; 11(3):357–374.

Fleming SM, Daw ND. Self-evaluation of decision-making: A general Bayesian framework for metacognitive computation. Psychological review. 2017; 124(1):91.

Fleming SM, Dolan RJ. In: Fleming SM, Frith CD, editors. The Neural Basis of Metacognitive Ability Berlin, Heidelberg: Springer Berlin Heidelberg; 2014. p. 245–265. https://doi.org/10.1007/978-3-642-45190-4_11, doi: 10.1007/978-3-642-45190-4_11.

Freedy A, DeVisser E, Weltman G, Coeyman N. Measurement of trust in human-robot collaboration. In: 2007 International symposium on collaborative technologies and systems Ieee; 2007. p. 106–114.

Friston K. The free-energy principle: a unified brain theory? Nature reviews neuroscience. 2010; 11(2):127–138.

Fu C, Yao X, Yang X, Zheng L, Li J, Wang Y. Trust Game Database: Behavioral and EEG Data From Two Trust Games. Frontiers in Psychology. 2019; 10. https://www.frontiersin.org/journals/psychology/articles/10.3389/fpsyg.2019.02656/full, doi: 10.3389/fpsyg.2019.02656, publisher: Frontiers.

Gilboa A, Marlatte H. Neurobiology of schemas and schema-mediated memory. Trends in cognitive sciences. 2017; 21(8):618–631.

Haring KS, Phillips E, Lazzara EH, Ullman D, Baker AL, Keebler JR. Applying the swift trust model to human-robot teaming. In: Trust in human-robot interaction Elsevier; 2021.p. 407–427.

Hauser TU, Iannaccone R, Stämpfli P, Drechsler R, Brandeis D, Walitza S, Brem S. The feedback-related negativity (FRN) revisited: new insights into the localization, meaning and network organization. Neuroimage. 2014; 84:159–168.

Jessup RK, Busemeyer JR, Brown JW. Error effects in anterior cingulate cortex reverse when error likelihood is high. Journal of Neuroscience. 2010; 30(9):3467–3472.

Kelly SP, O’Connell RG. Internal and external influences on the rate of sensory evidence accumulation in the human brain. Journal of neuroscience. 2013; 33(50):19434–19441.

Koban L, Pourtois G. Brain systems underlying the affective and social monitoring of actions: an integrative review. Neuroscience & Biobehavioral Reviews. 2014; 46:71–84.

Koban L, Pourtois G, Vocat R, Vuilleumier P. When your errors make me lose or win: event-related potentials to observed errors of cooperators and competitors. Social Neuroscience. 2010; 5(4):360–374.

Krueger F, McCabe K, Moll J, Kriegeskorte N, Zahn R, Strenziok M, Heinecke A, Grafman J. Neural correlates of trust. Proceedings of the National Academy of Sciences. 2007; 104(50):20084–20089.

Lebreton M, Abitbol R, Daunizeau J, Pessiglione M. Automatic integration of confidence in the brain valuation signal. Nature neuroscience. 2015; 18(8):1159–1167.

Li M, Tang D, Pan W, Zhang Y, Lu J, Li H. The influence of social status and promise levels in trust games: An Event-Related Potential (ERP) study. Cognitive, Affective, & Behavioral Neuroscience. 2025; 25(3):708–726. https://doi.org/10.3758/s13415-024-01259-9, doi: 10.3758/s13415-024-01259-9.

Loughnane GM, Newman DP, Bellgrove MA, Lalor EC, Kelly SP, O’Connell RG. Target selection signals influence perceptual decisions by modulating the onset and rate of evidence accumulation. Current Biology. 2016; 26(4):496–502.

Mathys CD, Lomakina EI, Daunizeau J, Iglesias S, Brodersen KH, Friston KJ, Stephan KE. Uncertainty in perception and the Hierarchical Gaussian Filter. Frontiers in human neuroscience. 2014; 8:825.

Mayer RC, Davis JH, Schoorman FD. An integrative model of organizational trust. Academy of management review. 1995; 20(3):709–734.

van der Meer J, Hoyte P, Roeder L, Bruza P, Modeling the quantum-like dynamics of human reliability ratings in Human-AI interactions by interaction dependent Hamiltonians; 2025. https://arxiv.org/abs/2504.13918.

Nassar MR, Wilson RC, Heasly B, Gold JI. An approximately Bayesian delta-rule model explains the dynamics of belief updating in a changing environment. Journal of Neuroscience. 2010; 30(37):12366–12378.

Nieuwenhuis S, Aston-Jones G, Cohen JD. Decision making, the P3, and the locus coeruleus–norepinephrine system. Psychological bulletin. 2005; 131(4):510.

Nightingale SJ, Farid H. AI-synthesized faces are indistinguishable from real faces and more trustworthy. Proceedings of the National Academy of Sciences. 2022; 119(8):e2120481119.

O’connell RG, Dockree PM, Kelly SP. A supramodal accumulation-to-bound signal that determines perceptual decisions in humans. Nature neuroscience. 2012; 15(12):1729–1735.

Oostenveld R, Praamstra P. The five percent electrode system for high-resolution EEG and ERP measurements. Clinical neurophysiology. 2001; 112(4):713–719.

Peirce J, Gray JR, Simpson S, MacAskill M, Höchenberger R, Sogo H, Kastman E, Lindeløv JK. PsychoPy2: Experiments in behavior made easy. Behavior research methods. 2019; 51:195–203.

Pernet C, Garrido MI, Gramfort A, Maurits N, Michel CM, Pang E, Salmelin R, Schoffelen JM, Valdes-Sosa PA, Puce A. Issues and recommendations from the OHBM COBIDAS MEEG committee for reproducible EEG and MEG research. Nature Neuroscience. 2020 Dec; 23(12):1473–1483. https://www.nature.com/articles/s41593-020-00709-0, doi: 10.1038/s41593-020-00709-0, publisher: Nature Publishing Group.

Pernet C, Rousselet G, Mas IS, Martinez R, Wilcox R, Delorme A. ElectroEncephaloGraphy robust statistical linear modelling using a single weight per trial. bioRxiv. 2022; https://www.biorxiv.org/content/early/2022/01/17/2021.04.27.441629, doi: 10.1101/2021.04.27.441629.

Pernet CR, Chauveau N, Gaspar C, Rousselet GA. LIMO EEG: a toolbox for hierarchical LInear MOdeling of ElectroEncephaloGraphic data. Computational intelligence and neuroscience. 2011; 2011(1):831409.

Pernet CR, Latinus M, Nichols TE, Rousselet GA. Cluster-based computational methods for mass univariate analyses of event-related brain potentials/fields: A simulation study. Journal of neuroscience methods. 2015; 250:85–93.

Pion-Tonachini L, Kreutz-Delgado K, Makeig S. ICLabel: An automated electroencephalographic independent component classifier, dataset, and website. NeuroImage. 2019; 198:181–197.

Polich J. Updating P300: an integrative theory of P3a and P3b. Clinical neurophysiology. 2007; 118(10):2128–2148.

Polich J. Neuropsychology of P300. In: Luck SJ, Kappenman ES, editors. The Oxford handbook of event-related potential components Oxford university press; 2012.p. p. 159–88.

Pothos EM, Busemeyer JR. Can quantum probability provide a new direction for cognitive modeling? Behavioral and brain sciences. 2013; 36(3):255–274.

Roeder L, Hoyte P, van der Meer J, Fell L, Johnston P, Kerr G, Bruza P. A Quantum Model of Trust Calibration in Human–AI Interactions. Entropy. 2023; 25(9):1362.

van Schie HT, Mars RB, Coles MG, Bekkering H. Modulation of activity in medial frontal and motor cortices during error observation. Nature neuroscience. 2004; 7(5):549–554.

Shadlen MN, Kiani R. Decision making as a window on cognition. Neuron. 2013; 80(3):791–806.

Simpson JA. Psychological foundations of trust. Current directions in psychological science. 2007; 16(5):264–268.

Somon B, Campagne A, Delorme A, Berberian B. Human or not human? Performance monitoring ERPs during human agent and machine supervision. Neuroimage. 2019; 186:266–277.

Ullsperger M, Fischer AG, Nigbur R, Endrass T. Neural mechanisms and temporal dynamics of performance monitoring. Trends in cognitive sciences. 2014; 18(5):259–267.

Wang Y, Zhang Z, Jing Y, Valadez EA, Simons RF. How do we trust strangers? The neural correlates of decision making and outcome evaluation of generalized trust. Social Cognitive and Affective Neuroscience. 2016; 11(10):1666–1676. https://academic.oup.com/scan/article/11/10/1666/2413988, doi: 10.1093/scan/nsw079.

