## Supplementary Material PDF for "Distinct brain mechanisms support trust violations, belief integration, and bias in human-AI teams" (from another version of this preprint: version 1, 2025-08-19)

### Accuracy of human judgements of images

The following figure (S1) shows the accuracy levels of the image judgments for all participants.

Averaged across participants, our participants were 50% (± 6%) accurate in judging whether a face was fake or real. Note that some participants seem to do very well (>60% accuracy for participant 019), while others are doing quite poor (<40% for participant 032).

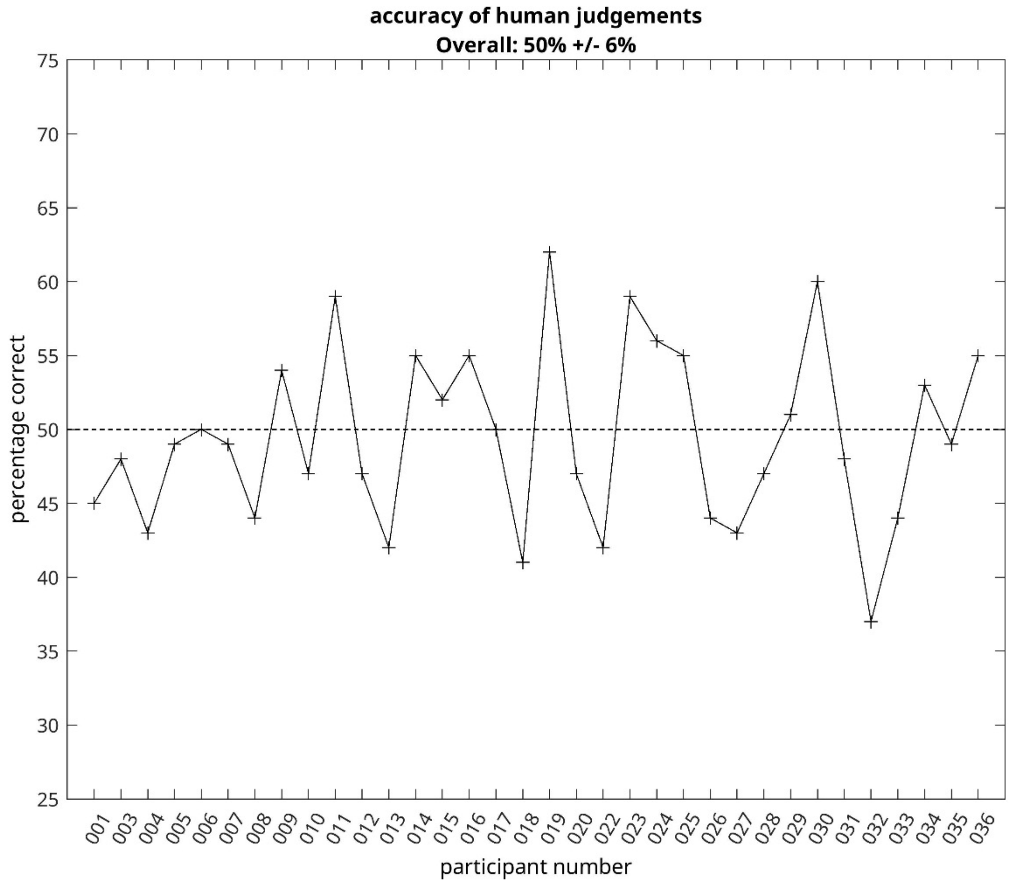

**Figure S1**. Accuracy of image judgments for all participants.

### Beta estimates

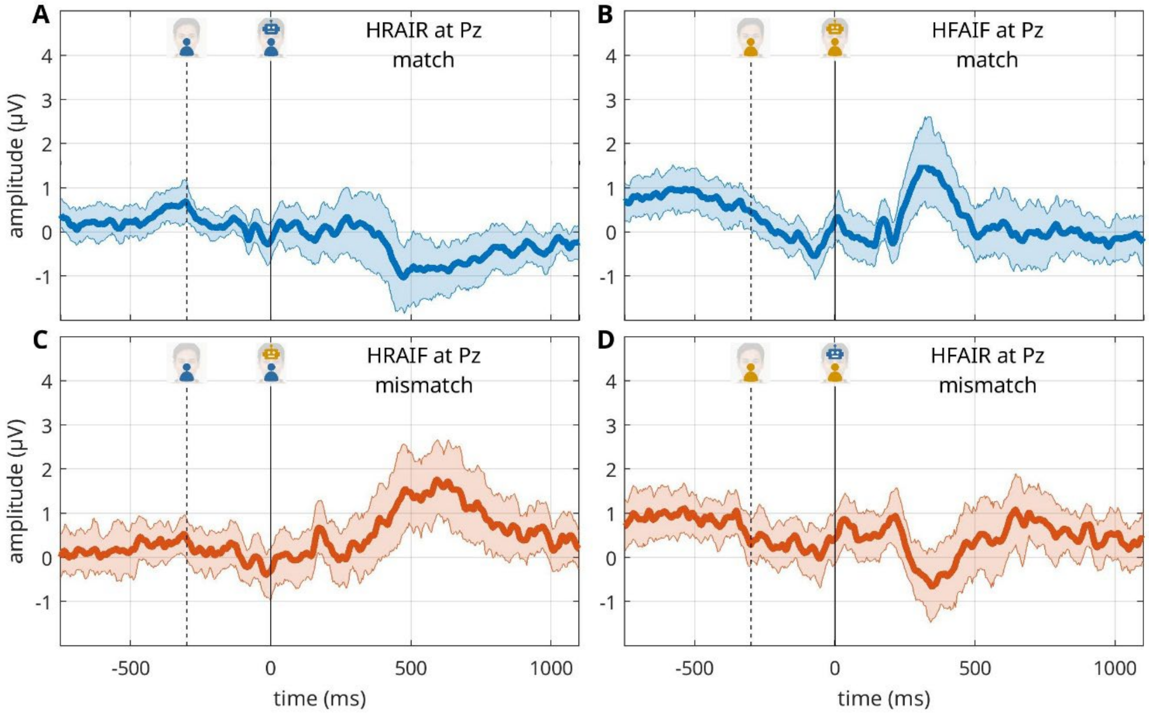

**Figure S2. Time series of betas** at Pz time-locked to the event when the AI classification icon (robot icon) is displayed on screen (time 0 ms) in the mismatch (red) and match (blue) conditions. At -300 ms participants made their judgment (fake or real key press). The match condition contains HRAIR and HFAIF, the mismatch condition contains HRAIF and HFAIR. Shaded areas denote 95% confidence intervals. HRAIR, Human-Real & AI-Real; HFAIF, Human-Fake & AI-Fake; HRAIF, Human-Real & AI-Fake; HFAIR, Human-Fake & AI-Real.

### Post-hoc t-test of AI classification (mismatch vs match)

After spatiotemporal clustering at an alpha level of 5%, mismatched AI feedback differed from matched AI feedback at one cluster from -184 ms to 1016 ms spanning all 64 electrodes (av. t = 0.64, av. p = 0.012). The channel-wise maximum statistic is at 296 ms.

To investigate the different temporal and topological patterns that are visible in Figure 3 and Figure S3, we also performed a spatiotemporal clustering correction at an alpha level of 1%. This revealed three clusters: 1) from 176 ms to 392 ms spanning 30 channels (av. t = 0.63, av. p = 0.002) with a channel-wise maximum statistic at 296 ms and a topological distribution centered around TP10. This cluster corresponds to the N2 ERP component. 2) from 392 ms to 552 ms spanning 59 channels (av. t = 3.22, av. p = 0.002) with a channel-wise maximum statistic at 460 ms and a topological distribution centered around FC1. This cluster corresponds to the P3a ERP component. 3) from 496 ms to 772 ms spanning 47 channels (av. t = 1.09, av. p = 0.003) with a channel-wise maximum statistic at 700 ms and a topological distribution centered around P8. This cluster corresponds to the P3b ERP component.

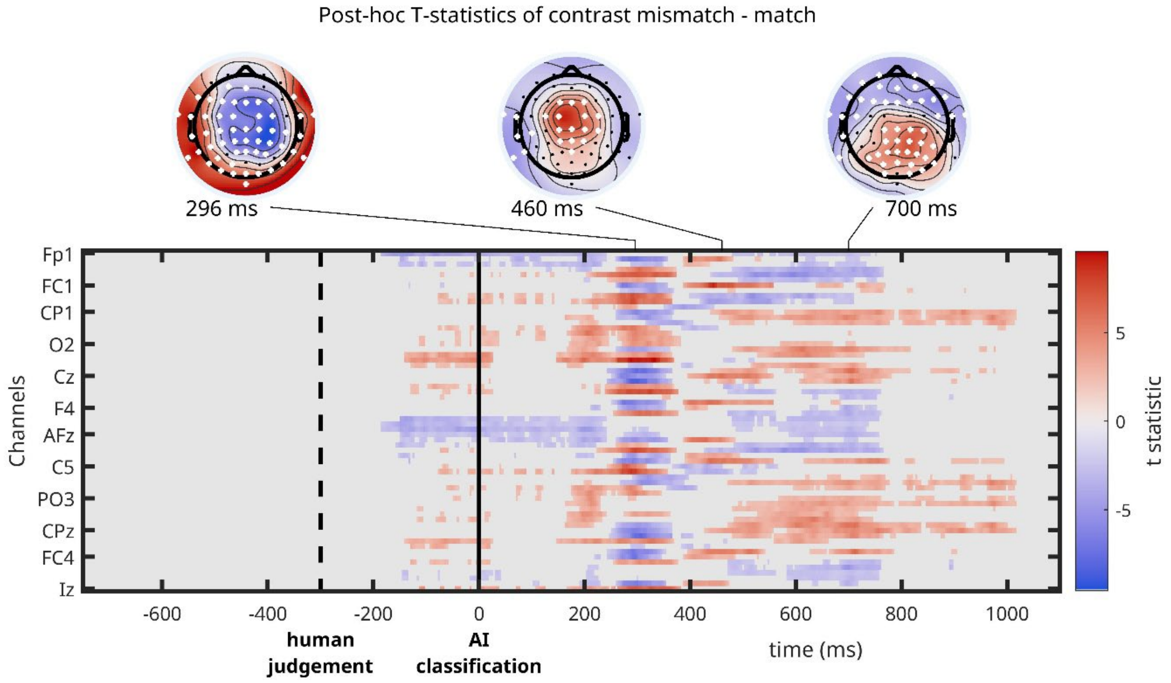

**Figure S3.** Results from post-hoc paired *t*-test for the AI-classification-locked betas comparing the mismatch with the match condition shown as heat maps indicating significant regions and latencies (red, positive *t*-values; blue, negative *t*-values). Channels are sorted according to their relative proximity to each other. Topological distributions are presented at three latencies (296, 460 and 700 ms after AI feedback). Results are corrected with spatiotemporal clustering at an alpha level of 5%.

### Post-hoc t-test for interaction of AI classification (mismatch vs match) and human judgment of images (fake vs real)

After spatiotemporal clustering at an alpha level of 5%, the human judgment type (fake or real) impacted processing of the AI classification (mismatched, matched) differently at one cluster: from 256 ms to 796 ms spanning all 64 channels (av. t = -0.65, av. p = 0.011). The channel-wise maximum statistic is at 484 ms.

After spatiotemporal clustering at an alpha level of 1%, we identified two main clusters: 1) from 312 ms to 404 ms spanning 20 channels (av. t = -1.52, av. p = 0.003) with a channel-wise maximum statistic at 360 ms and a topological distribution centered around P2. This cluster coincides with the N2 ERP period. 2) from 420 ms to 640 ms spanning 26 channels (av. t = -1.74, av. p = 0.002) with a channel-wise maximum statistic at 484 ms and a topological distribution centered around P3. This cluster coincides with the P3b ERP period.

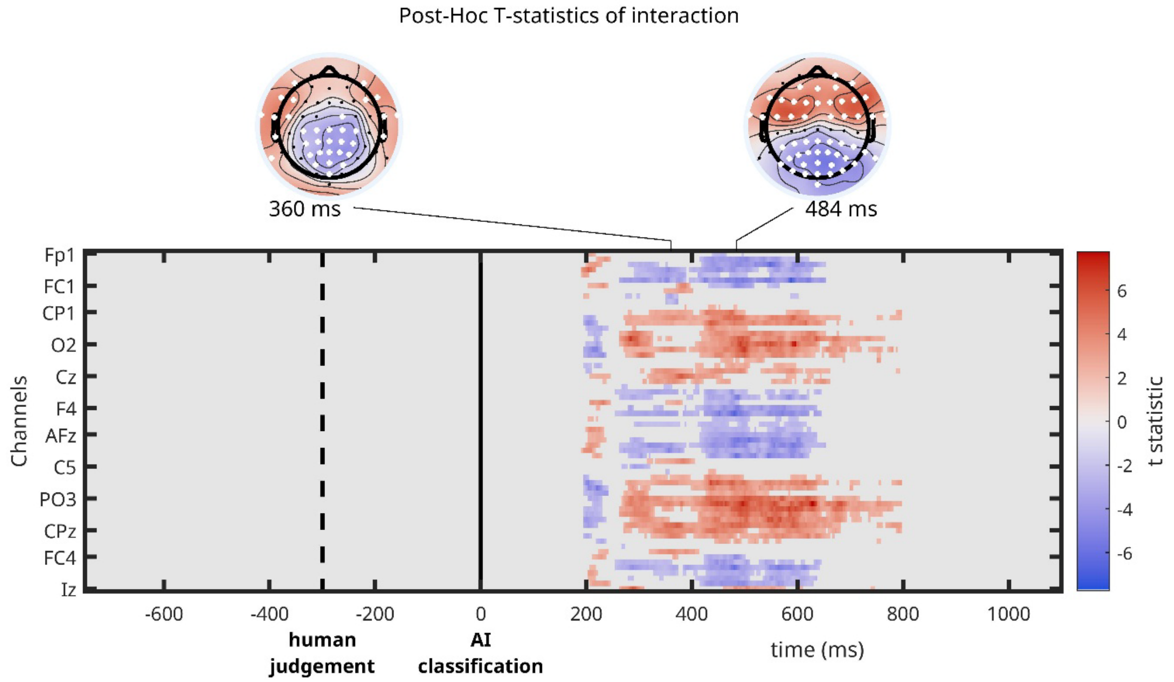

**Figure S4.** Results from post-hoc paired *t*-test of the interaction effect of AI classification (mismatch, match) after a human image judgment of fake versus real, shown as heat maps indicating significant regions and latencies (red, positive *t*-values; blue, negative *t*-values). Channels are sorted according to their relative proximity to each other. Topological distributions are presented at two latencies (360 and 484 ms after AI feedback). Results corrected with spatiotemporal clustering at an alpha level of 5%.

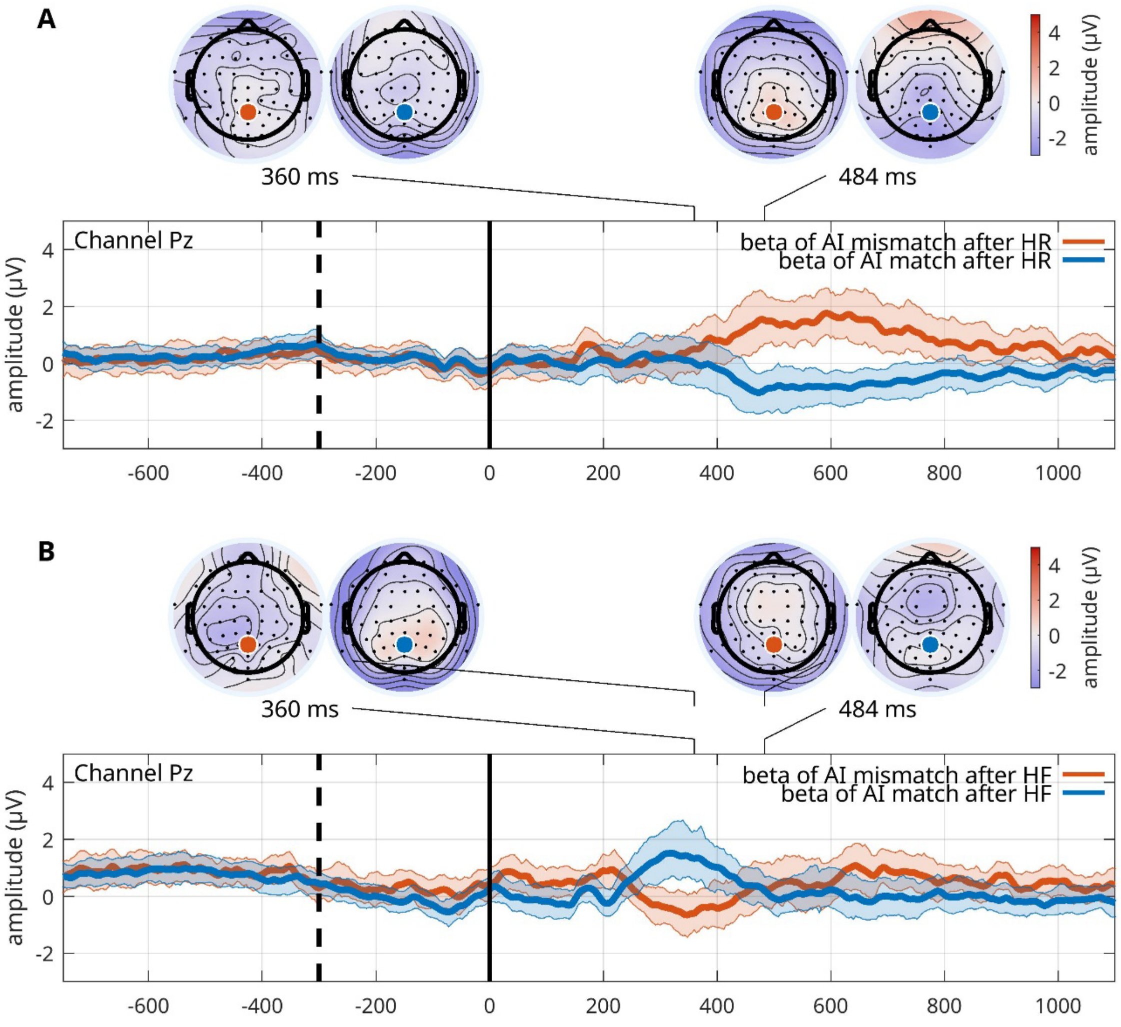

**Figure S5.** Time series and topologies of betas of interaction effect at channel Pz time-locked to the event of AI classification (time 0 ms). **A.** When humans judged an image as real, the beta time series of the mismatch (red) and match (blue) conditions, with topologies at two latencies (360 ms, 484 ms) with Pz highlighted in red (mismatch), blue (match). **B.** Shows the equivalent to panel A when humans judged an image as fake. Beta time series are presented as the 20% trimmed mean, shaded areas denote the 95% Bayesian Highest Density Interval. HF, Human Fake; HR, Human Real.

Figure S6 (below) illustrates the interaction effect we found. It highlights that the processing of the AI classification (mismatched or matched) is dependent on the preceding human judgment of the image (real or fake).

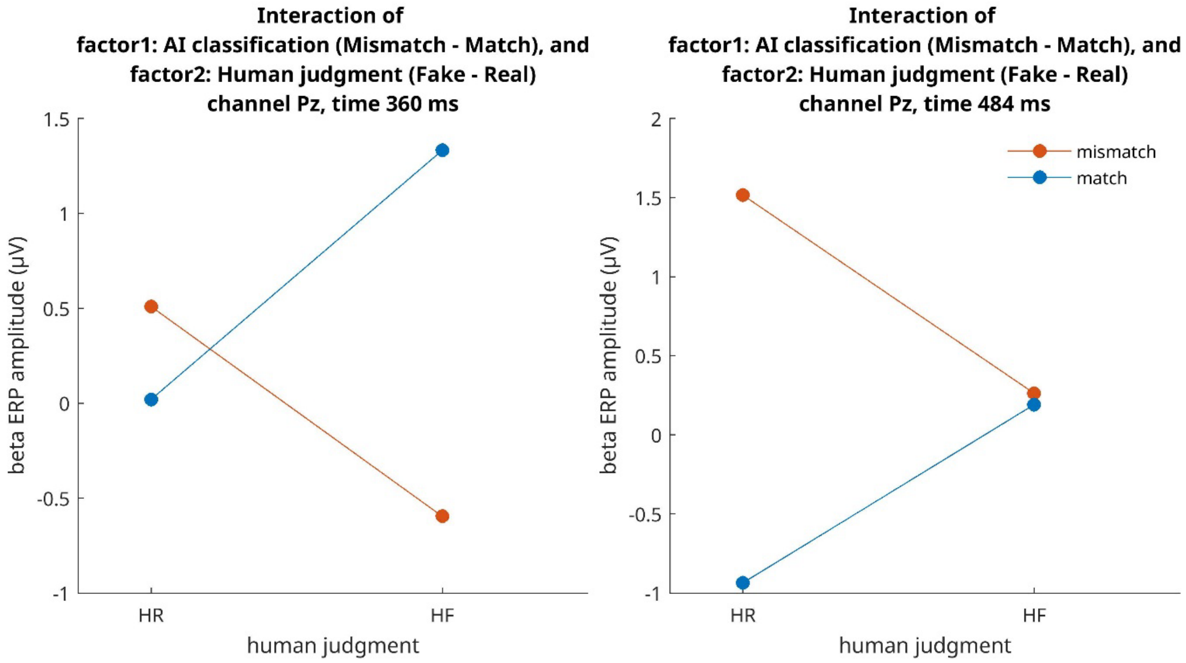

**Figure S6. Illustration of interaction effect at two latencies.** ERP beta amplitude at 360 ms (left) and at 484 ms (right) at channel Pz when human judged an image as real (HR) and when human judged an image as fake (HF) with mismatched (red) and matched AI feedback (blue). HF, Human Fake; HR, Human Real.

### Post-hoc t-test for human judgment of images (fake vs real)

Both spatiotemporal clustering at an alpha level of 5% and at an alpha level of 1% identified one cluster. At an alpha level of 1%, human judgment of fake differed from a human judgment of real at one cluster from -724 ms to -420 ms spanning 16 channels (av. t = 3.74, av. p = 0.003). The channel-wise maximum statistic is at -508 ms centered around P4.

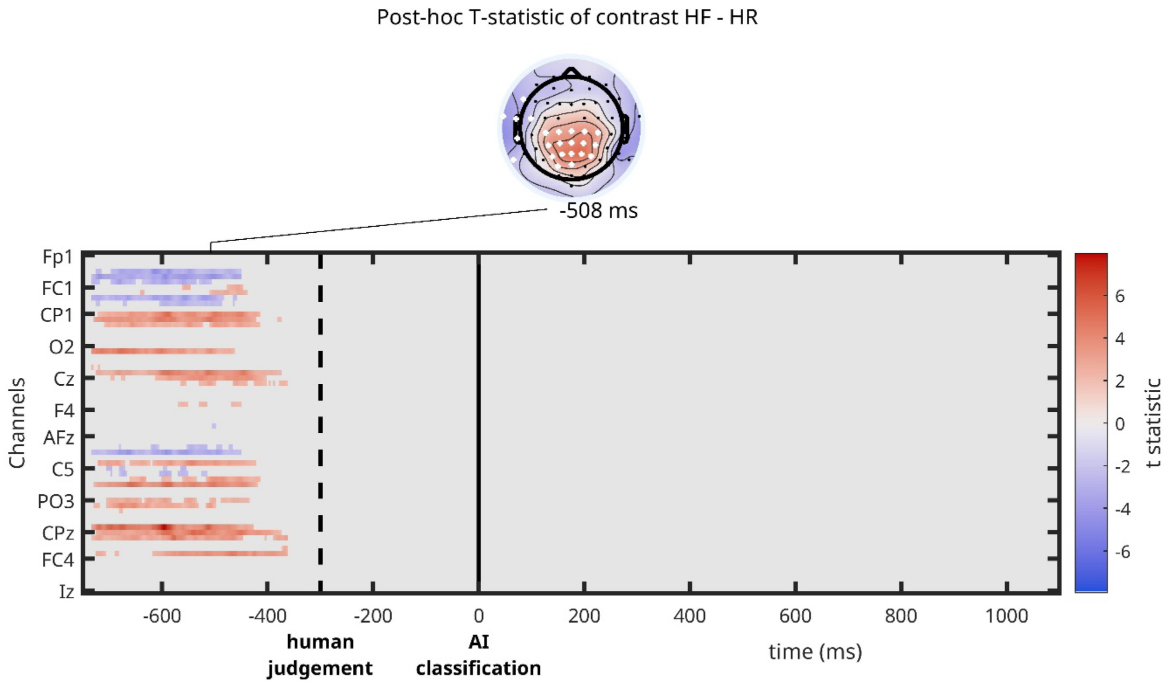

**Figure S7.** Results from post-hoc paired *t*-test for the AI-classification-locked betas comparing the human judgement conditions (fake versus real) shown as heat maps indicating significant regions and latencies (red, positive *t*-values; blue, negative *t*-values). Channels are sorted according to their relative proximity to each other. Topological distributions are presented at one latency (-508 ms before AI classification). Results are corrected with spatiotemporal clustering at an alpha level of 5%. HF, Human Fake; HR, Human Real.

### Reliability ratings for all participants

The ratings vary widely across participants and are highly personal. For most participants the ratings are centered around the individual mean rating. The span of the y-axis (the rating) is from -25% to +25% centered around the participant's mean which is reported in the figure. This is to be able to compare the variability of the ratings across participants.

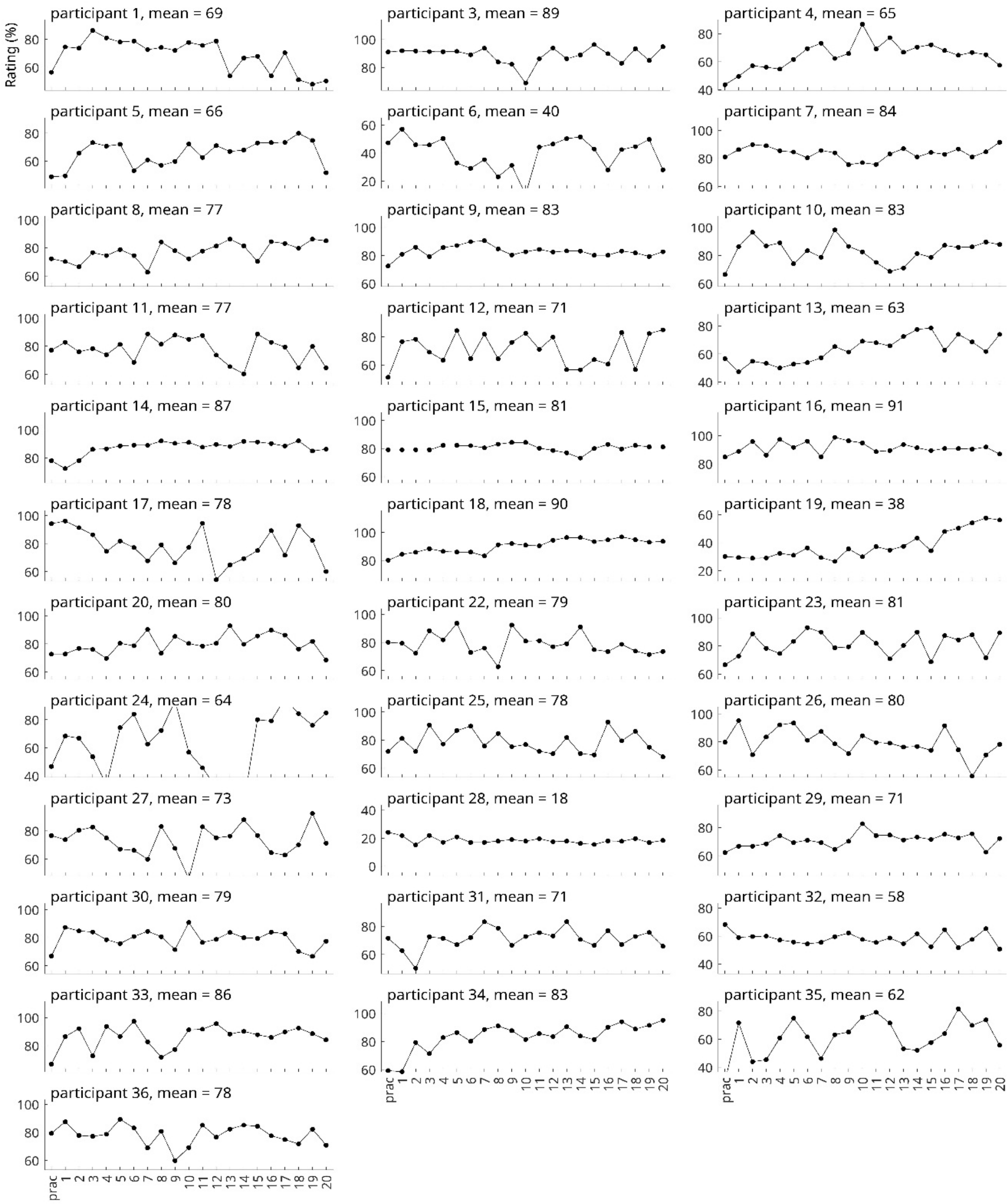

**Figure S8**. Reliability ratings for each individual participant. Each panel shows the reliability ratings in % for each block and participant (x axis depicts the block number, y axis shows the % reliability rating); the mean rating (in %) across blocks for each participant is reported at the top of each panel.

### Post-hoc t-test for co-variation of ERP amplitudes and changes in reliability ratings

We show post-hoc t-tests considering HRAIR trials in Figure S9; HFAIF in Figure S10; HRAIF in Figure S11, and HFAIR in Figure S12. Only Figure S9 is controlled for false positives by cluster-wise correction; Figures S10, S11 and S12 are not controlled for false positives and are shown for display purposes only.

For HRAIR (Figure S9), the t-test at an alpha level of 5% revealed three significant clusters: 1) from -632 to -444 ms spanning 20 channels (av. t = 0.04, av. p = 0.016) with a channel-wise maximum statistic at -564 ms and a topological distribution centered around P7; 2) from -144 to -24 ms spanning 16 channels (av. t = -2.73, av. p = 0.018) with a channel-wise maximum statistic at -112 ms and a topological distribution centered around Fp1; 3) from 172 to 268 ms with 48 channels (av. t = -2.84, av. p = 0.017) with a channel-wise maximum statistic at 184 ms and a topological distribution centered around Fp1.

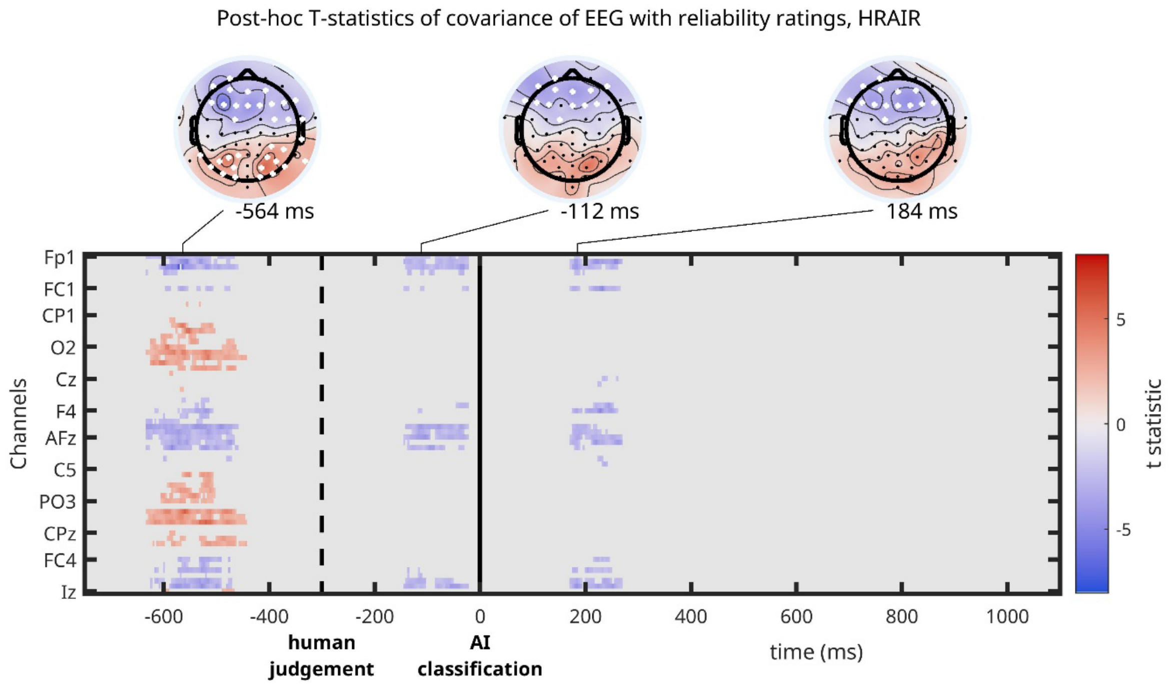

**Figure S9**. Results from post-hoc paired *t*-test for the AI-classification-locked betas checking for co-variation of ERP amplitude and changes in reliability ratings for Human-Real-AI-Real (HRAIR) trials (human judgment was real, AI feedback was matched), shown as heat maps indicating significant regions and latencies (red, positive *t*-values; blue, negative *t*-values). Channels are sorted according to their relative proximity to each other. Topological distributions are presented at three latencies (-564 ms, -112 ms and 184 ms before/after AI feedback). Electrodes belonging to significant clusters are marked white; non-significant electrodes are marked black. Results corrected with spatiotemporal clustering at an alpha level of 5%.

Post-hoc T-statistics of covariance of EEG with reliability ratings, HFAIF

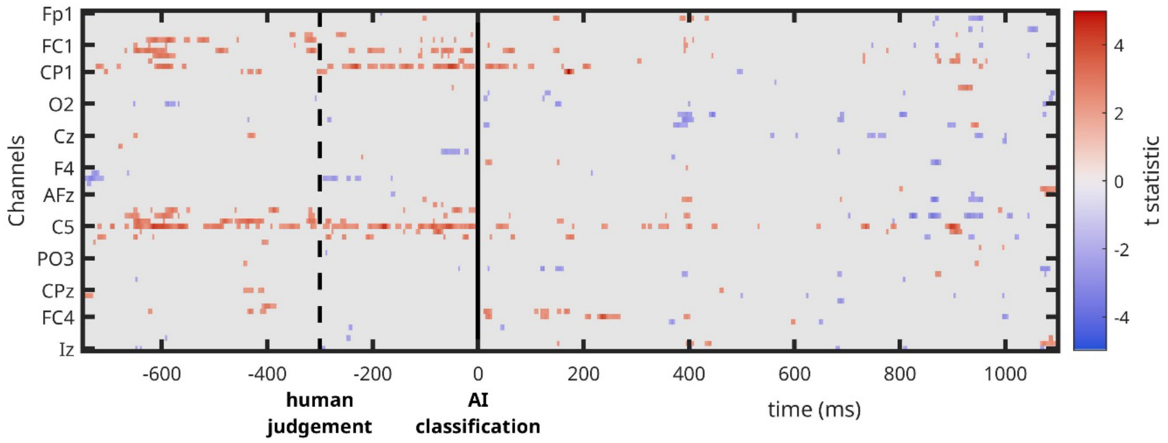

**Figure S10.** Results from post-hoc paired t-test for the AI-classification-locked betas checking for co-variation of ERP amplitude and changes in reliability ratings for Human-Fake-AI-Fake (HFAIF) trials (human judgment was fake, AI feedback was matched), shown as heat maps indicating significant regions and latencies (red, positive *t*-values; blue, negative *t*-values). Channels are sorted according to their relative proximity to each other. Results are not corrected with spatiotemporal clustering. The t-test did not reveal any significant clusters.

Post-hoc T-statistics of covariance of EEG with reliability ratings, HRAIF

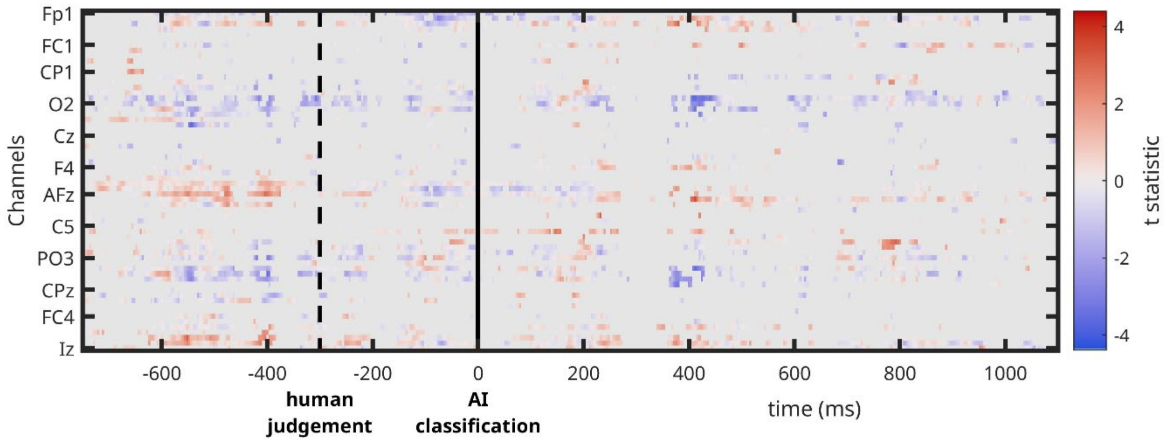

**Figure S11.** Results from post-hoc paired *t*-test for the AI-classification-locked betas checking for co-variation of ERP amplitude and changes in reliability ratings for Human-Real-AI-Fake (HRAIF) trials (human judgment was real, AI feedback was mismatched), shown as heat maps indicating significant regions and latencies (red, positive *t*-values; blue, negative *t*-values). Channels are sorted according to their relative proximity to each other. Results not corrected with spatiotemporal clustering. The t-test did not reveal any significant clusters.

Post-hoc T-statistics of covariance of EEG with reliability ratings, HFAIR

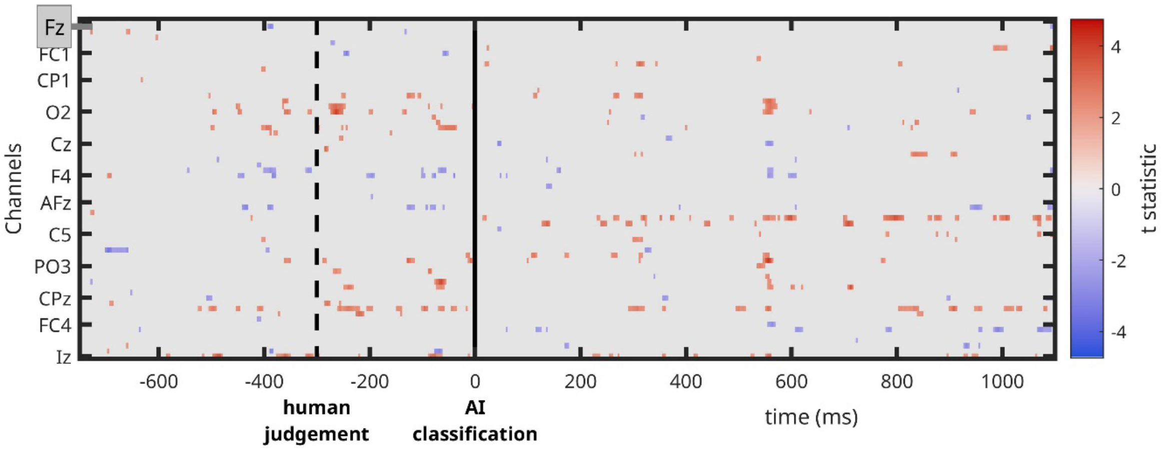

**Figure S12.** Results from post-hoc paired *t*-test for the AI-classification-locked betas checking for co-variation of ERP amplitude and changes in reliability ratings for Human-Fake-AI-Real (HFAIR) trials (human judgment was fake, AI feedback was mismatched), shown as heat maps indicating significant regions and latencies (red, positive *t*-values; blue, negative *t*-values). Channels are sorted according to their relative proximity to each other. Results are not corrected with spatiotemporal clustering. The t-test did not reveal any significant clusters.

### Correlations between ERP amplitudes and changes in reliability ratings per block for all participants

Most participants (n = 25) showed a positive correlation between ERP amplitude and change in reliability rating at channel Fz and latency -520 ms.

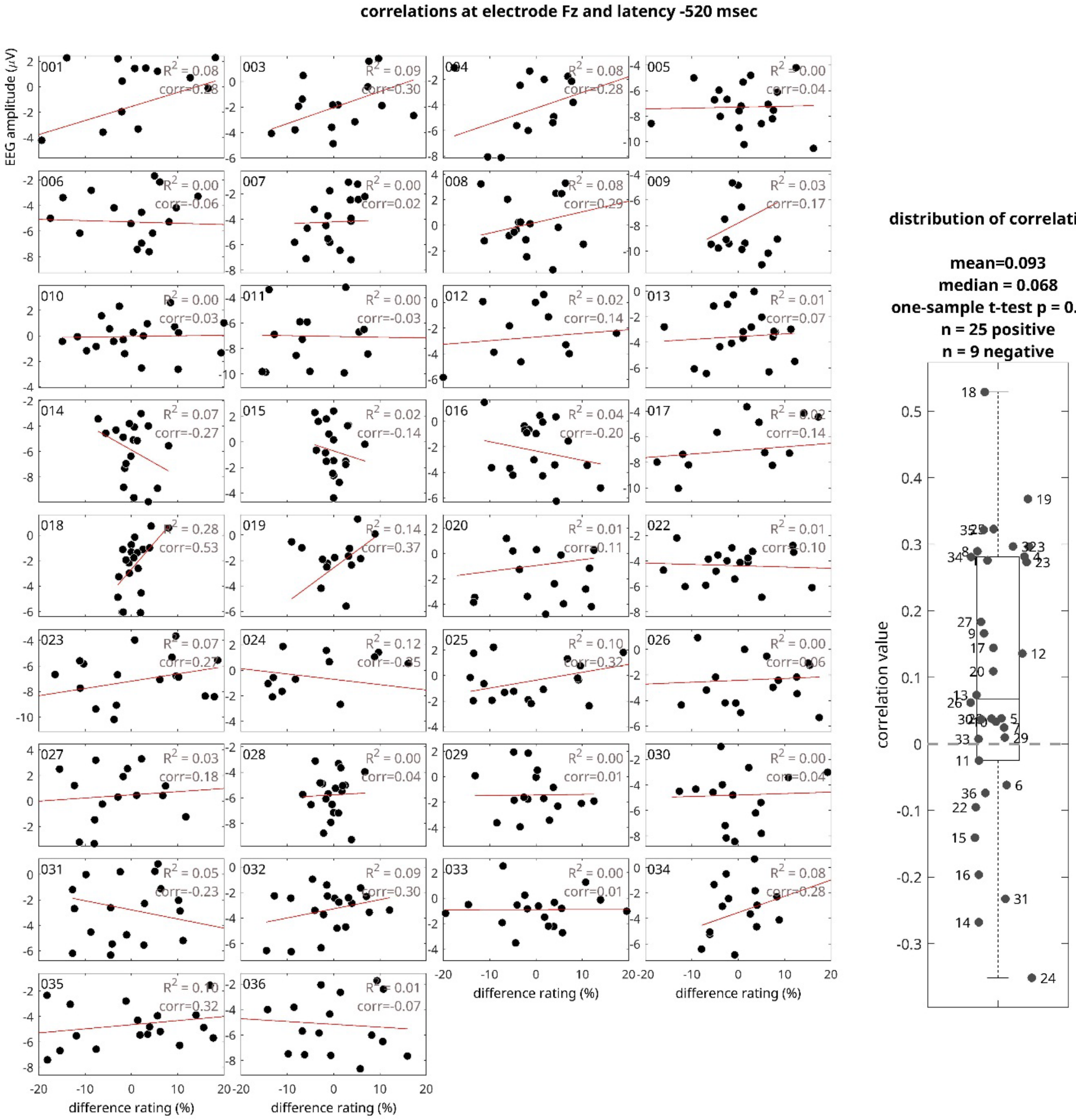

**Figure S13.** Correlations between ERP amplitude and difference reliability rating per block for each participant (left). Each panel shows the correlation between the difference reliability rating per block (post-pre) and the mean ERP amplitude per block (averaged across all HRAIR trials per block) at Fz at -520 ms for each participant (each black dot shows estimate of one block); regression lines of best fit are displayed in red. Boxplot (right) shows the distribution of correlation coefficients across participants (showing the median, 25th and 75th percentile, minimum and maximum).

### Post-hoc t-test for co-variation of ERP amplitudes (mismatch-match beta) and human response bias

The t-test revealed four significant clusters: 1) from -96 ms to 292 ms spanning 41 electrodes (av. t = -1.09, av. p = 0.013); the channel-wise maximum statistic is at 120 ms, with topological distributions centered around PO3. 2) from 464 ms to 748 ms spanning 18 electrodes (av. t = -2.79, av. p = 0.014); the channel-wise maximum statistic is at 548 ms, with topological distributions centered around AF4. 3) from 492 ms to 784 ms spanning 32 electrodes over frontal and parietal regions (av. t = 2.54, av. p = 0.02); the channel-wise maximum statistic is at 640 ms, with topological distributions centered around P5. 4) from 808 ms to 884 ms spanning 18 electrodes (av. t = 2.50, av. p = 0.022); the channel-wise maximum statistic is at 836 ms, with topological distributions centered around CP5.

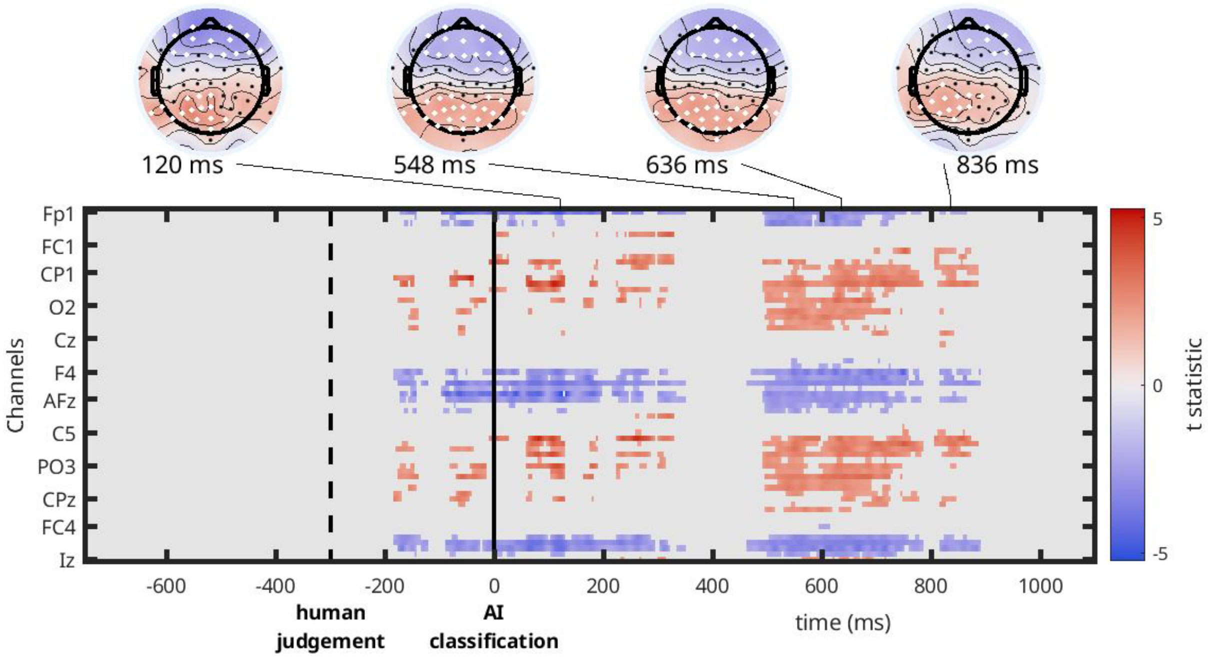

**Figure S14.** Results from post-hoc paired *t*-test for the AI-classification-locked betas checking for co-variation of human response bias and ERP amplitude (mismatch-match beta difference wave), shown as heat maps indicating significant regions and latencies (red, positive *t*-values; blue, negative *t*-values). Channels are sorted according to their relative proximity to each other. Topological distributions are presented at four latencies (120 ms, 548 ms, 636 ms and 836 ms after AI classification). Electrodes belonging to significant clusters are marked white; non-significant electrodes are marked black. Results corrected with spatiotemporal clustering at an alpha level of 5%.

### Regression results for co-variation of ERP amplitudes and human response bias during mismatched AI classification trials

This section outlines the results of the regression between response bias and mismatch beta estimates presented in supplementary Figure S15.

Three main significant clusters were found across similar time ranges. The first cluster spans 9 channels and time points from -188 to -100 ms (av. F = 6.61, av. p=0.022). The maximum F value across channels was at -136 ms with a topological distribution centered around channel POz. The second cluster spans 18 channels and time points ranging from -188 to 184 ms (av. F = 6.99, av. p = 0.019). The maximum F-value across channels was at 32 ms with a topological distribution centered around channel AF4. The third cluster spans 24 channels and time points ranging from -92 to 132 ms (av. F = 6.71, av. p = 0.021). The maximum F-value across channels was at 64 ms with a topological distribution around channel POz.

We plot the topological distributions at these three latencies in Figure S15. Additionally, we also plot the topography at 120 ms for ease of comparison with results in Figure 7.

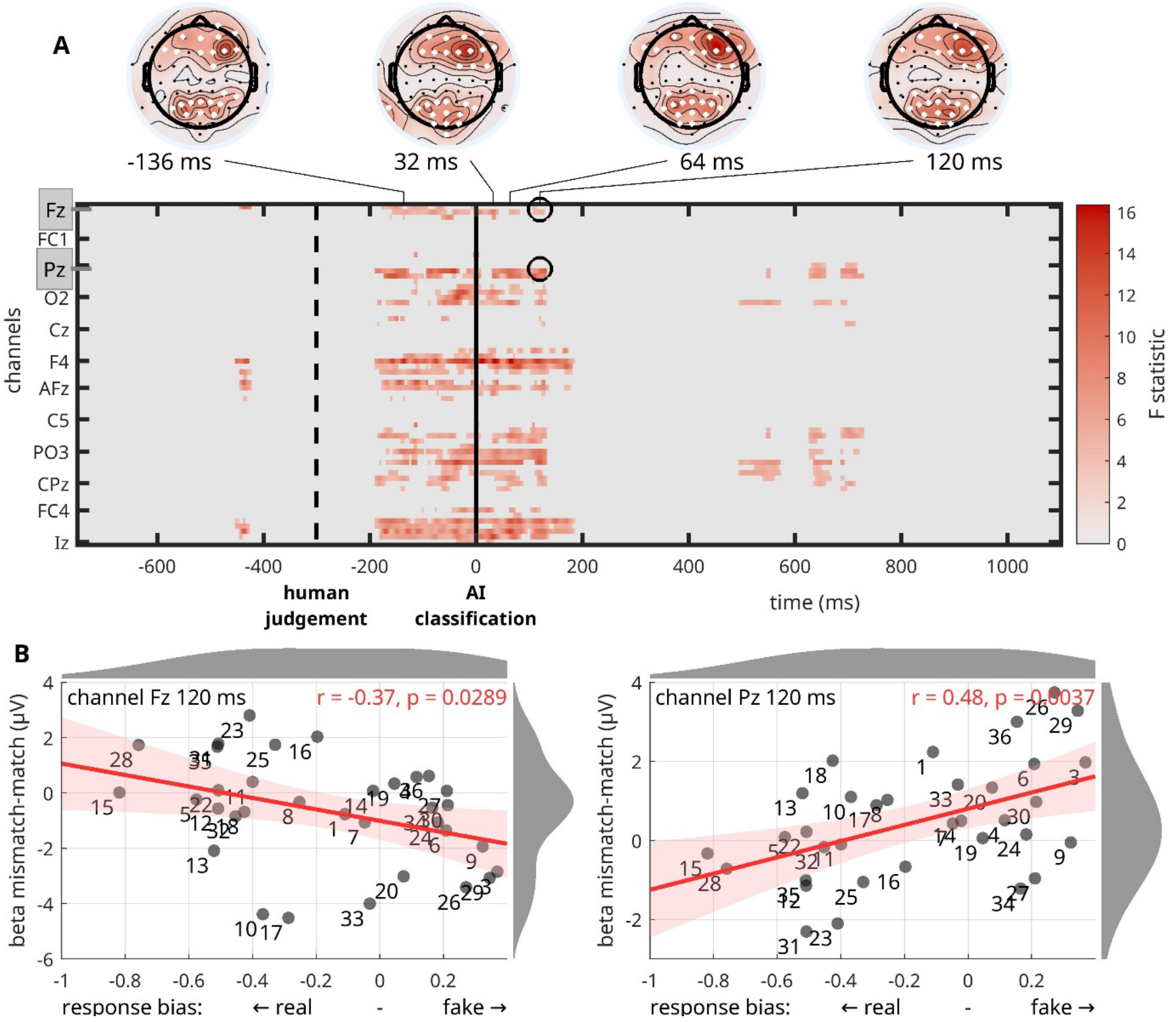

**Figure S15.** Regression results of human response bias for judging images as fake or real and ERP amplitude for the AI mismatching the human judgment. **A.** F-statistic results from linear regression for the AI-classification-locked betas checking for correlations of human bias for judging images as real or fake and ERP amplitude (beta) of the mismatch condition. Results are shown as heat map indicating significant F-values in red over channels and time points. Channels Fz and Pz are highlighted for further investigation in panel B. Topological distributions are presented at four latencies (-136 ms, 32 ms, 64 ms and 120 ms). Electrodes belonging to significant clusters are marked white; non-significant electrodes are marked black. Results are corrected with spatiotemporal clustering at an alpha level of 5%. **B.** Correlations between bias for judging an image as fake and the amplitude of the mismatch beta at channel Fz and Pz at 120 ms. Gray dots represent participants, red line shows regression across participants, red shaded area the 95% CI. X-axis: values > 0 indicate a bias for fake, < 0 for real. Gray shaded areas at the edge of panels show the distribution of the response bias across participants (x-axis direction) and the distribution of the mismatch beta amplitude (y-axis direction).

### Regression results for co-variation of ERP amplitudes and human response bias during matched AI classification trials

This section outlines the results of the regression between response bias and match beta estimates presented in supplementary Figure S16.

Three main significant clusters were found. The first cluster spans 12 channels and time points ranging from 224 to 392 ms (av. F = 7.45, av. p = 0.017). The maximum F-value across channels was at 320 ms with a topological distribution centered around channel T7. The second (and largest) cluster spans 56 channels and time points ranging from 480 to 1028 ms (av. F = 7.16, av. p = 0.018). The maximum F-value across channels was at 628 ms with a topological distribution at both frontal and parietal regions. The third cluster spans 9 channels and time points from 808 to 892 ms (av. F = 5.81, av. p=0.025). The maximum F-value across channels was at 864 ms with a topological distribution centered around channel Fp2.

To be consistent with the results presented in Figure 7, we also plot the regressions with their coefficients at 636 ms for Fz and Pz in supplementary figure S16B; this time point is within the time ranges of the second cluster.

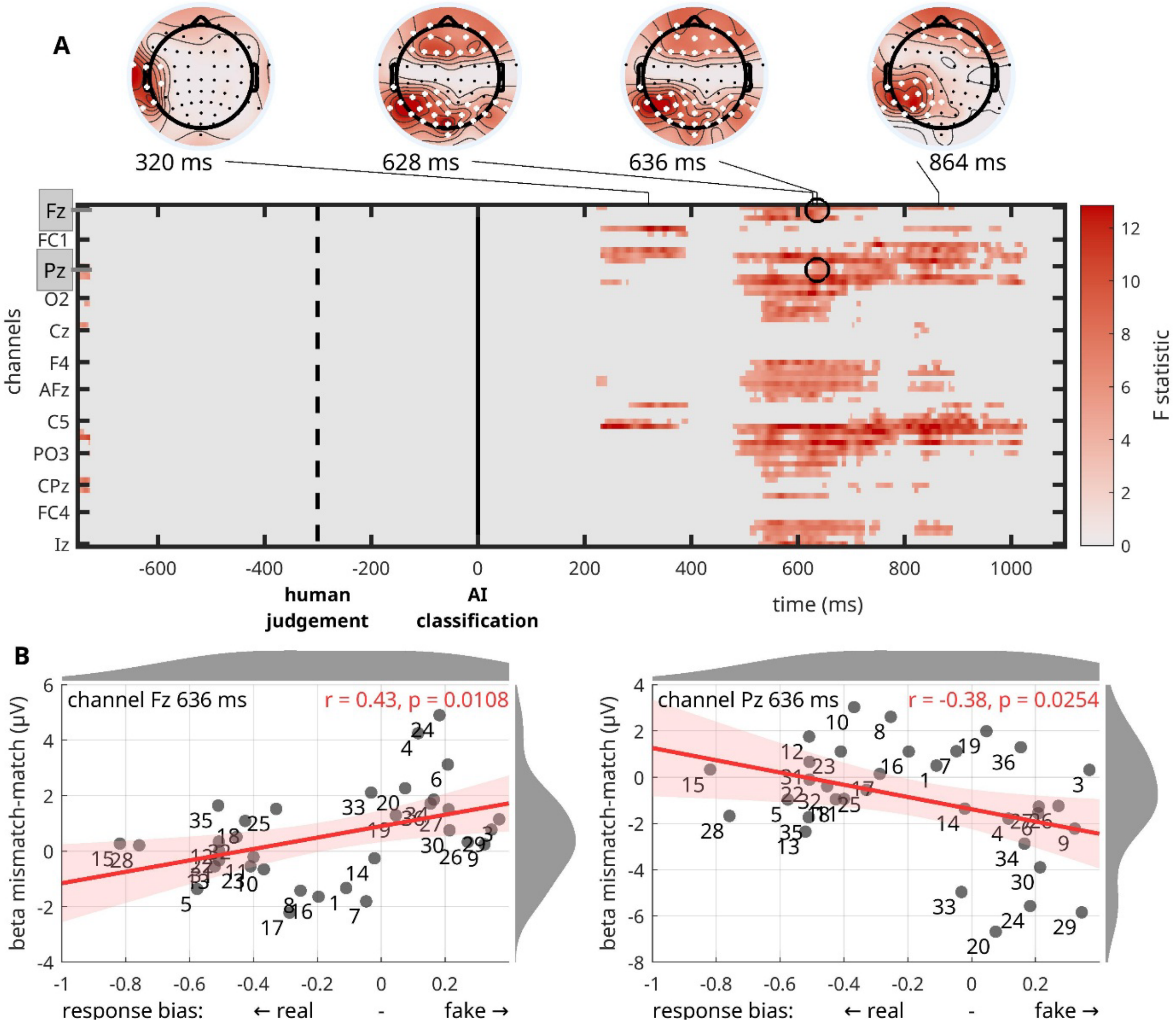

**Figure S16.** Regression results of human response bias for judging images as fake or real and ERP amplitude for the AI matching the human judgment. **A.** F-statistic results from linear regression for the AI-classification-locked betas checking for correlations of human bias for judging images as real or fake and ERP amplitude (beta) of the match condition. Results are shown as heat map indicating significant F-values in red over channels and time points. Channels Fz and Pz are highlighted for further investigation in panel B. Topological distributions are presented at four latencies (320 ms, 628 ms, 636 ms and 864 ms). Electrodes belonging to significant clusters are marked white; non-significant electrodes are marked black. Results are corrected with spatiotemporal clustering at an alpha level of 5%. **B.** Correlations between bias for judging an image as fake and the amplitude of the match beta at channel Fz and Pz at 636 ms. Gray dots represent participants, red line shows regression across participants, red shaded area the 95% CI. X-axis: values > 0 indicate a bias for fake, < 0 for real. Gray shaded areas at the edge of panels show the distribution of the response bias across participants (x-axis direction) and the distribution of the match beta amplitude (y-axis direction).
